# Clusters, Fingers, and Singles: A Mechanical Landscape of Tumor Invasion

**DOI:** 10.1101/2025.11.20.689434

**Authors:** Sheriff Akeeb, Adam I Marcus, Yi Jiang

## Abstract

Collective invasion is a key mechanism by which tumors disseminate and metastasize, involving coordinated migration of heterogeneous cell populations. Experimental studies have identified specialized leader and follower cells that work together during this process, but the biophysical rules governing their interaction remain unclear. We present a mechanistic, cell-based computational model using the Cellular Potts framework to investigate how heterotypic adhesion, leader motility, and follower proliferation jointly shape invasion. Leader–follower tumors were simulated across 13,310 parameter sets, and invasion was quantified by invasive and infiltrative areas, finger-like protrusions, solitary defectors, and detached clusters. From these simulations, we identified four distinct invasion phenotypes: non-invasive, bulk collective, single-cell, and multimodal. Multimodal invasion—the coexistence of cohesive strands, solitary cells, and small clusters—emerged as the most prevalent phenotype, particularly under moderate adhesion, high motility, and intermediate proliferation. Proliferation exhibited nonlinear effects: moderate division reinforced cohesive invasion, whereas excessive growth destabilized tumor architecture. Mapping outcomes across the parameter space revealed sharp transitions between invasion modes, underscoring trade-offs between adhesion and motility in shaping invasion complexity. Our results show that hybrid invasion behaviors, previously considered rare, arise robustly from simple mechanical rules and are favored in a broad region of the parameter space. This framework reconciles binary models of invasion with experimental observations of heterogeneity, providing predictive insights into how modulating adhesion, motility, or proliferation can restrict metastatic spread.

**Author summary:** When cancer cells leave a primary tumor, they do not just invade as lone cells but also as cohesive fingers and small cell clusters, depending on how strongly cells stick together, how hard some cells pull, and how fast the population grows. We built a simple, physics-based computational model in which a small number of ‘leader’ cells pull on the more proliferative ‘followers.’ By systematically varying three key properties, cell-cell adhesion, leader motility, and follower proliferation, we generated a large simulation of invasion patterns and grouped them into four invasion modes. The resulting phase diagram shows where tumors grow as compact masses (no invasion), or invade as protruding “fingers,” as single cells, or in a mixed multimodal fashion with fingers, clusters, and single cells.

Surprisingly, this mixed state was the most common outcome, arising when adhesion is moderate and leader motility is strong. This map provides intuition and practical rules for thinking about how changes in adhesion or motility might reorganize invasion, such as shifting cluster-forming fronts toward more compact, less dissemination-prone configurations.

## Introduction

Metastasis, the leading cause of cancer-related mortality, often begins with local invasion, where tumor cells breach the basement membrane and infiltrate surrounding tissues. This process involves diverse cellular strategies that enable migration through complex microenvironments and ultimately facilitate colonization of distant sites [1, 2]. A growing body of evidence distinguishes between two primary modes of invasion: solitary migration, where individual cells disseminate independently, and collective invasion, where groups of cells migrate in a coordinated, cohesive manner [3–5].

Solitary invasion is frequently associated with epithelial-to-mesenchymal transition (EMT), a process characterized by the loss of epithelial adhesion, upregulation of mesenchymal markers, and enhanced motility and invasiveness [5, 6]. In contrast, collective invasion preserves cell–cell junctions and spatial organization, relying on mechanical coordination, intercellular signaling, and polarity among neighboring cells [7]. This mode of invasion is increasingly recognized across multiple carcinoma types, including breast [8], colorectal [9], and squamous cell cancers [10], and is associated with enhanced metastatic efficiency, immune evasion, and resistance to apoptosis [11].

A signature of collective invasion is the emergence of functional heterogeneity among migrating tumor cells, particularly the distinction between leader and follower subpopulations [12–15]. Leader cells, typically localized at the invasive front, exhibit high motility, proteolytic activity, and directional sensing, actively remodeling the extracellular matrix (ECM). Follower cells, in contrast, remain less motile but contribute to structural cohesion and proliferation within the invading collective. This division of labor is reminiscent of migratory behaviors observed in development, such as neural crest migration and border cell clusters [16].

Recent experimental advances, including the Spatiotemporal Genomic and Cellular Analysis (SaGA) platform, have enabled the molecular profiling of leader and follower cells, revealing transcriptional and phenotypic signatures that confirm their distinct roles in invasion [12, 17]. These findings underscore the importance of spatial heterogeneity within tumors and raise critical questions about how biophysical properties such as cell–cell adhesion, motility, and proliferation give rise to diverse invasion behaviors.

Despite these insights, most computational models of cancer invasion simplify the invasion spectrum into binary categories, collective or solitary, and often neglect the spatial, mechanical, and phenotypic dynamics observed in real tumors [18–22]. In particular, few models explicitly represent leader–follower heterogeneity or explore how interactions among adhesion, motility, and proliferation shape invasion phenotypes. As a result, those models often fail to capture the complex, hybrid behaviors increasingly observed in aggressive cancers, such as the coexistence of collective strands, solitary cells, and detached clusters within a single tumor front [23].

To address this gap, we develop a biophysical, cell-based computational framework that explicitly incorporates leader–follower dynamics within a two-dimensional spatial domain. Using the Cellular Potts Model (CPM) [24, 25], we systematically vary key adhesion, motility, and proliferation parameters to map the emergent patterns of invasion behaviors. We simulate over 13,000 unique conditions to explore the emergent invasion behaviors.

We hypothesize that the interplay among adhesion, motility, and proliferation governs transitions between distinct invasion modes, and that multimodal invasion, characterized by simultaneous collective strands, single-cell dispersal, and detached clusters, arises robustly under specific biophysical regimes. Our goal is to construct a high-resolution phenotype map that links local cell-level properties to tissue-scale invasion strategies, providing both mechanistic insight and predictive capacity.

By integrating mechanistic modeling with data-driven classification, our study advances the theoretical understanding of collective invasion and leader–follower coordination, offering a quantitative framework to interpret tumor heterogeneity and potentially inform therapeutic strategies aimed at disrupting metastatic dissemination.

## Results

### Distinct Invasion Metric Sensitivities to Adhesion, Motility, and Proliferation

To dissect how cellular biophysical properties shape emergent tumor invasion dynamics, we systematically varied leader–follower contact energy (*J*_*lf*_), leader migration coefficient (λ), and follower proliferative probability (PP) across a three-dimensional parameter space encompassing 13.310 simulations (1,331 unique parameter combinations with n = 10 replicates each). Each simulation ran for 700 MCS (approximately 36 h of biological time), and we quantified five invasion metrics at endpoint: invasive area (expansion of the connected tumor mass), infiltrative area (total spatial footprint including dispersed cells), number of finger-like protrusions, single invading cells (singles), and detached clusters. These metrics capture distinct aspects of tumor spatial organization and collectively define invasion phenotypes.

Invasive area, measuring the expansion of the connected tumor mass, was maximized under intermediate adhesion (*J*_*lf*_ ≈ 0 to 2) and elevated migration (λ ≥15), particularly when paired with moderate-to-high proliferation (PP ≥ 0.5) (S2 Fig). Strong adhesion (*J*_*lf*_ ≤ − 2) yielded mean invasive areas of 5420 px^2^, compared to 10 314 px^2^ under intermediate adhesion (− 2 < *J*_*lf*_ ≤ 2) and 6697 px^2^ under weak adhesion (*J*_*lf*_ > 2). Migration exhibited a stronger effect: low migration (λ < 10) produced mean invasive areas of 2906 px^2^, whereas high migration (λ ≥20) yielded 12 180 px^2^, representing a 4.2-fold increase. Strong adhesion prevented structural remodeling, while very weak adhesion reduced cohesion, both limiting expansion. These trends suggest a synergy between motility and proliferation: leader-driven pulling must be reinforced by proliferative replenishment in the tumor core to sustain coherent expansion.

In contrast, the infiltrative area, which captures the full spatial footprint including dispersed single cells and clusters, was dictated primarily by migration. It increased monotonically with λ, rising from 3579 px^2^ at low migration to 30 592 px^2^ at high migration (8.5-fold increase), and peaked under weak adhesion (*J*_*lf*_ > 2) at 29 345 px^2^ (S3 Fig). Thus, directional motility alone is sufficient for extensive tumor spread, although it does not necessarily lead to cohesive growth.

Finger counts peaked at intermediate adhesion (*J*_*lf*_ ≈ 0 to 2) with mean = 7.11 fingers, and strong migration (λ ≥ 20) with mean = 9.74 fingers (S5 Fig). Strong adhesion (*J*_*lf*_ ≤ −2) restricted protrusive branching (mean = 4.03 fingers), while weak adhesion (*J*_*lf*_ > 2) fragmented collectives into singles or clusters (mean = 6.11 fingers). This non-monotonic dependence on adhesion is consistent with experimental studies showing that optimal intercellular adhesion promotes dynamic collective invasion [26, 27]. Proliferation had a negligible influence on finger formation (Pearson r = −0.000, p = 0.985). The number of singles increased sharply with λ and weak adhesion (*J*_*lf*_ > 2), reflecting a transition toward mesenchymal-like escape (S4 Fig). Weak adhesion produced mean = 175.79 singles compared to 1.53 under strong adhesion and 53.06 under intermediate adhesion. While such singles enable early dissemination, clinical evidence suggests that solitary circulating tumor cells frequently enter dormancy or undergo apoptosis, limiting their metastatic potential [28].

Detached clusters, defined as cohesive aggregates that separate from the core, formed in 28.3 % of all simulations (3763 out of 13 310) and were most prevalent under intermediate-to-weak adhesion (*J*_*lf*_ = 0 to 3) and high migration (λ ≥ 24) (S6 Fig). Intermediate adhesion produced a mean of 1.69 clusters per simulation, while weak adhesion yielded 1.78 clusters. High migration (λ ≥ 24) resulted in 70.1 % of simulations exhibiting clusters, with mean of 5.00 clusters when present. In this regime, adhesion is strong enough to preserve intercellular cohesion within the aggregate yet weak enough to allow detachment from the primary mass. Larger clusters tended to persist until the end of simulations (700 MCS), while smaller aggregates often dissolved earlier. Again, proliferation (PP) showed no consistent effect on cluster frequency or stability (Pearson r = − 0.002, p = 0.815). These observations parallel experimental findings that cluster-based dissemination is highly metastatic [29, 30].

Correlation-based sensitivity analysis (S7 Fig) corroborated these trends: migration showed the strongest correlations with invasive area (Pearson r = 0.70, Spearman ρ = 0.80), infiltrative area (Pearson r = 0.69, Spearman ρ = 0.80), and finger counts (Pearson r = 0.80, Spearman ρ = 0.81). Contact energy exhibited moderate-to-strong correlations with singles (Pearson r = 0.67, Spearman ρ = 0.71) and infiltrative area (Pearson r = 0.57, Spearman ρ = 0.50). Proliferation exhibited weak correlation across all metrics (Pearson and Spearman |r| < 0.03, all p > 0.05), reinforcing its limited role in determining invasion mode.

Together, these findings demonstrate leader cell motility and intercellular adhesion as the primary regulators of invasion architecture, with proliferation acting mainly as a scaling factor for tumor bulk. This functional decoupling underscores the central role of mechanical and migratory properties in shaping invasion phenotype and spatial spread.

### Phenotypic Phase-Space Mapping Reveals Prevalence of Multi-modal Invasion

We classify emergent invasion behaviors within our leader-follower framework into four biologically interpretable phenotypes: No Invasion, Single-Cell Invasion, Bulk Invasion, and Multimodal Invasion. Phenotype labels were assigned by a rule-based classifier applied to multiple invasion metrics, including invasive area, infiltrative spread, finger counts, single cells, and detached clusters (see Model and Methods). This approach provides an objective mapping of complex invasion behaviors across the parameter landscape.

These categories align with the framework proposed by Friedl *et al*. [5], which organizes invasion behavior along a spectrum from single-cell migration to collective streaming. Our Single-Cell Invasion corresponds to mesenchymal or amoeboidal motility of leader cells under weak intercellular contacts. Bulk Invasion reflects coordinated, strand-like migration with preserved junctional integrity. No Invasion represents non-migratory, growth-dominant behavior. Finally, Multimodal Invasion shows the richest dynamics, with simultaneous formation of collective fingers, singles, and detached clusters, a phenotype increasingly observed in aggressive clinical tumors [29, 31] but rarely captured in computational models. Fig 1 and S8 Fig illustrate how these phenotypes emerge over time.

**Fig 1.**
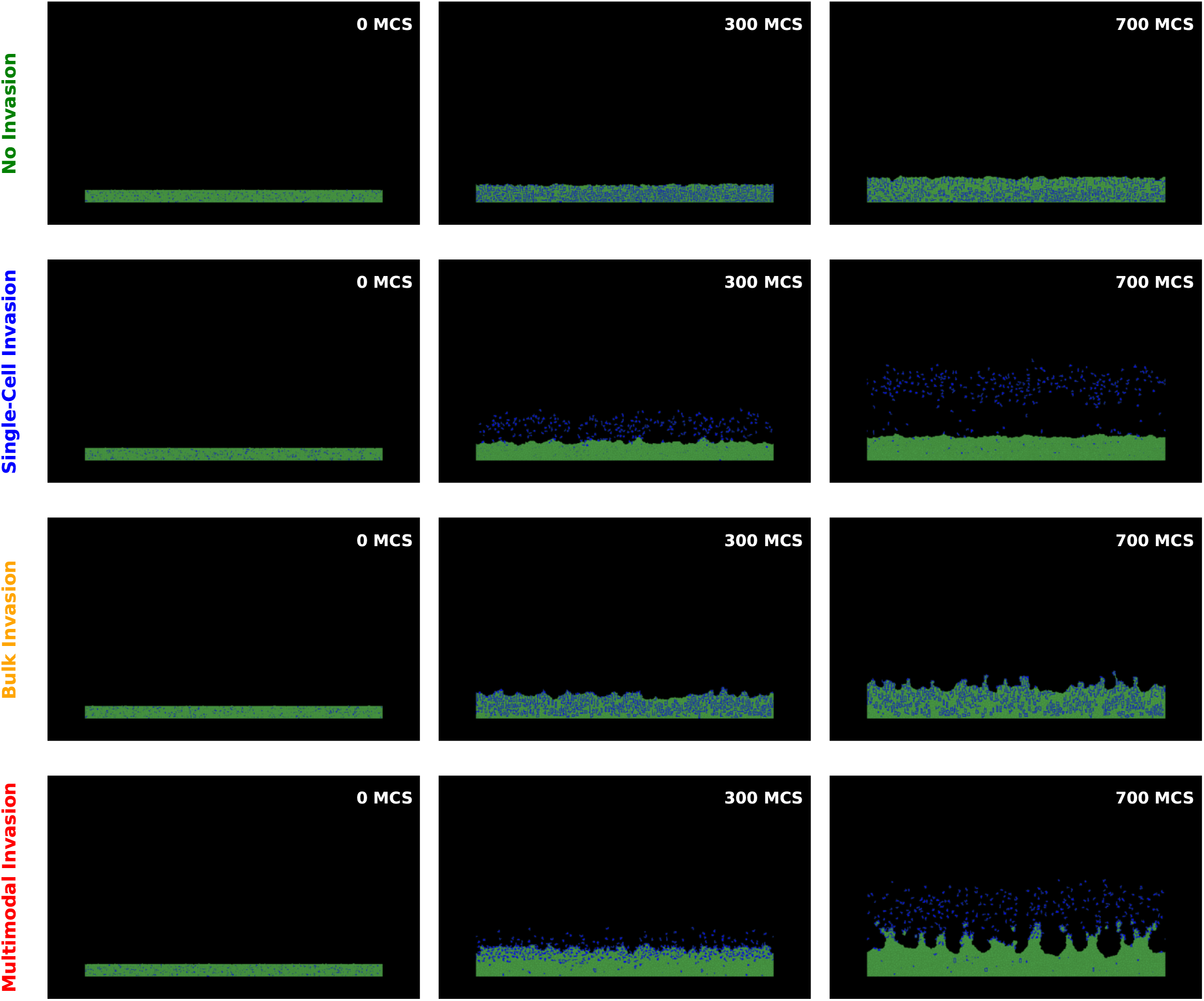
Temporal evolution of tumor morphologies across invasion phenotypes. Rows show representative simulations for the four phenotypes, no invasion, single cell invasion, bulk invasion, and multimodal invasion, while columns show time points (0, 300, and 700 MCS, corresponding to 0, 18, and 32 hours). Leaders are blue and followers are green. No invasion (top row) remains compact, illustrating regimes of strong adhesion, negligible leader motility, and low proliferation. Single-cell invasion (second row) emerges under weak adhesion and high leader motility, where leader cells detach and migrate individually. Bulk invasion (third row) features cohesive, finger-like protrusions that preserve junctional integrity, supported by high motility and proliferative replenishment. Multimodal invasion (bottom row) exhibits coexisting fingers, dispersed singles, and detached clusters, under intermediate adhesion, high leader motility, and moderate proliferation. See S1 video for full spatiotemporal dynamics.

We visualized phenotype assignments in the 3D parameter space defined by leader–follower contact energy (*J*_*lf*_), leader migratory coefficient (λ), and follower proliferative probability (PP). The resulting phase map (Fig 2A) reveals well-delineated phenotypic regions separated by sharp, nonlinear transitions as one or more control parameters vary. As summarized in S1 Table, No Invasion phenotype (green) occupies approximately 22% of the parameter space primarily at low migration (λ < 10) and moderate to strong adhesion (*J*_*lf*_ ≤ 2), where leader cells either lack migratory drive or remain tightly bound to follower cells, preventing outward expansion. Single-Cell Invasion (blue) appears in a narrow but distinct band (~ 1% of parameter space) at weak adhesion (*J*_*lf*_ > 3), low to intermediate migration (4 ≤ λ ≤ 12), and minimal proliferation, where leaders detach and migrate independently with limited follower engagement. Bulk Collective Invasion (orange) emerges in approximately 23% of parameter space under strong adhesion (*J*_*lf*_ ≪ 0), moderate migration (λ ≈ 10 − 20), and moderate proliferation (0.1 < PP < 0.7), producing cohesive strands resembling the leader-follower chains observed in NSCLC spheroids [12]. In contrast, Multimodal Invasion (gold) spans a broad range (~ 54% of parameter space) of intermediate to weak adhesion (− 1 ≤ *J*_*lf*_ ≤ 5), high migration (λ > 15), and moderate to high proliferation (PP > 0.3), representing the most prevalent invasion outcome in our simulation.

**Fig 2.**
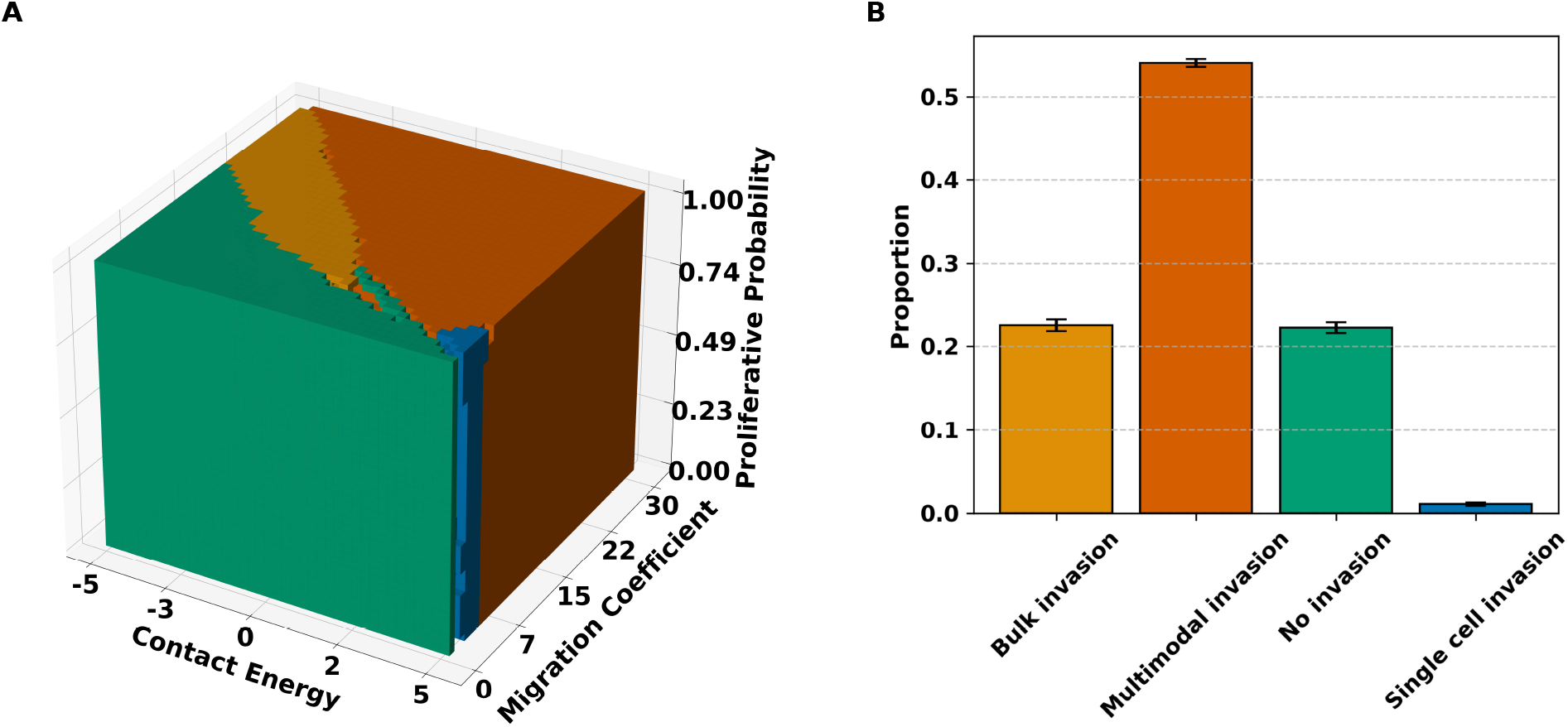
3D phase space of invasion phenotypes. (**A**) Multidimensional phenotype map over leader–follower contact energy (*J*_*lf*_), leader migration strength (*λ*), and follower proliferative probability (PP). Each voxel is assigned the most probable phenotype: green (No Invasion), blue (Single-Cell Invasion), orange (Bulk Invasion), or gold (Multimodal Invasion). Boundaries between regimes are sharp and nonlinear, and multimodal invasion occupies a large region characterized by moderate adhesion, strong migration, and intermediate-to-high proliferation. See S8 Fig for 2D cross-sections of the phase space. (**B**) The phenotype frequency distribution across 13, 310 simulations. Multimodal invasion is the most prevalent outcome, where bulk and no invasion occur under more restricted conditions, and single-cell invasion is comparatively rare. Error bars denote mean ± s.d. across 10 replicates. S4 Video shows the temporal progression, with early compact states giving way to increasing multimodal prevalence at later times.

Notably, multimodal invasion is the most prevalent outcome in our simulations (Fig 2B), with more than half of all parameter combinations favoring this heterogeneous phenotype. This suggests that heterogeneous invasion, characterized by coexisting finger-like protrusions, detached clusters, and isolated individual cells, is a robust and recurrent tumor behavior. This phenotype emerges most frequently under intermediate adhesion and high motility, consistent with histopathological and experimental observations in invasive carcinomas, where collective strands often coexist with solitary migratory cells and multicellular clusters [10, 29]. While specific tumor subtypes may differ in their parameter mappings, the phase structure suggests testable strategies for shifting tumors away from multimodal regions by jointly modulating adhesion and motility.

### Cluster Formation Peaks Under Intermediate Adhesion and Strong Migration

Multicellular clusters are potent mediators of metastatic spread, as circulating tumor cell (CTC) clusters demonstrate markedly higher metastatic efficiency than individual CTCs [11, 29, 30]. Clusters typically preserve leader–follower heterogeneity, wherein leader cells provide traction and directional persistence while followers maintain cohesion. These cellular collectives therefore represent a distinct invasion mode, distinct from purely single-cell escape or bulk-like advance.

To examine the mechanistic drivers of cluster detachment, we systematically evaluated simulation outcomes and computed cluster-related metrics at every 10 MCS, including cluster counts, size distributions, cellular composition, and trajectory persistence, across 13 310 simulations.

Cluster emergence depended strongly on the adhesion-motility balance. Heatmaps of cluster incidence (S9A, C Fig) showed a sharp increase under intermediate adhesion (*J*_*lf*_ ≈ 0–3) and strong motility (λ ≈ 24–30). These conditions define a mechanically permissive regime in which adhesion is low enough to permit detachment yet high enough to preserve intra-cluster cohesion, while high leader motility provides leader-driven traction for escape. By contrast, strong adhesion (*J*_*lf*_ < − 2) or low motility (λ < 15) suppressed cluster formation, confining cells to the primary tumor mass. Phase maps colored by cluster persistence (Fig 3, S11) further demonstrated that longer-lived clusters predominantly occurred in this intermediate-adhesion, high-migration window. Across the explored range, PP exerted at most a secondary influence on cluster incidence, though it can modulate size and longevity once clusters form.

**Fig 3.**
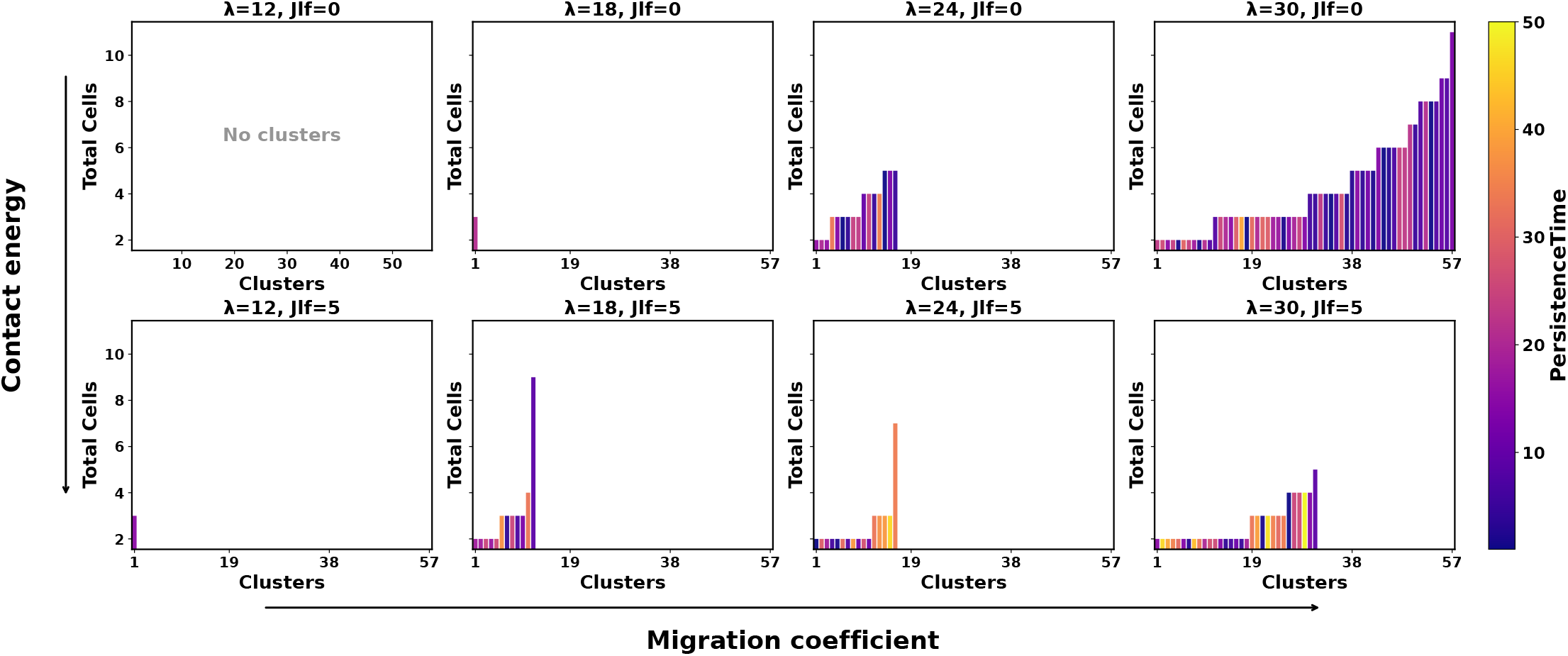
Representative cluster size distributions across migration and adhesion at fixed proliferation. Shown for *PP* = 0.9 at a subset of (*λ, J*_*lf*_) combinations chosen to illustrate regimes with and without cluster formation. (See S11 Fig for the full panel.) Each panel shows the distribution of cluster sizes (y-axis) versus the number of clusters observed (x-axis), pooled across *n* = 10 replicates and sampled every 10 MCS. Bars are colored by persistence time, with lighter shades indicating longer-lived clusters. Panels are arranged in the (*λ, J*_*lf*_) plane, with migration coefficient increasing from left to right and contact energy increasing from top to bottom, as indicated by the outer axis arrows. Panels labeled “No Clusters” indicate conditions under which no clusters formed. Across the representative conditions, strong-adhesion or low-motility regimes show few or no clusters, whereas intermediate adhesion (*J*_*lf*_ ≈ 0–2) combined with high motility (*λ* ≥ 24) yields numerous small-to-intermediate clusters and occasional large, persistent aggregates, capturing the heterogeneous outcomes characteristic of this parameter window.

The aggregate distribution of cluster sizes was strongly right-skewed or heavy-tailed (S9B Fig). Most clusters consisted of 4-8 cells, while rare aggregates exceeded 30 cells. The mean cluster size was approximately seven cells. Persistence time increased with both size and leader fraction. Small clusters typically dissipated within 10-20% of the total simulation time (70 MCS – 140 MCS), whereas larger, leader-enriched clusters often persisted until the final stages (500 MCS –700 MCS). The inset of S11 Fig highlights (λ = 30, *J*_*lf*_ = 2), where small clusters dominated but rare large aggregates persisted for the full duration of the simulation, reflecting heterogeneous outcomes characteristic of this regime.

Trajectory analyses showed that centroid displacement increased with cluster size and motility. At (*J*_*lf*_ = 1, λ = 30, PP = 0.9), for example, leader-rich clusters maintained persistent displacement over time, while smaller clusters exhibited short trajectories and frequent dissolution (S10 Fig). Velocity analysis confirmed that larger clusters and those enriched in leaders sustained higher speeds, underscoring the pivotal role of leaders in driving directional invasion. Clusters with lower leader fractions or smaller size displayed reduced velocity and shorter lifespans.

Cluster stability was reflected in cellular composition. Box plots of leader/follower counts per cluster (S9D Fig) revealed that leaders consistently outnumbered followers (medium 4 vs. ~ 3 per cluster), and the leader fraction distribution peaked at ~ 60% (S9E Fig). This enrichment indicates that clusters are not random aggregates but are leader-dominated collectives with robust composition. Larger clusters with higher leader content exhibited enhanced stability and longer persistence, while smaller or follower-rich clusters dissolved more readily.

Proliferation had a comparatively minor effect on cluster incidence and persistence across the parameter space. Varying PP altered the follower availability but did not qualitatively change the adhesion–motility trends or cluster stability. Fixing PP = 0.9 in parameter scans confirmed that proliferation primarily increased follower counts within clusters (S9D Fig, bottom) without eliminating leader dominance. This finding suggests that motility and adhesion, rather than proliferation, are the principal determinants of cluster detachment and longevity.

To further examine cluster dynamics, we tracked splitting and merging events. Splits (one cluster dividing into smaller aggregates or single cells) and merges (detached clusters rejoining another cluster or the tumor margin) were most frequent under intermediate adhesion and high motility, where mechanical interactions allowed partial cohesion and dynamic rearrangement. (S2 video) illustrates these events and corroborates the trajectories in S10 Fig.

These results align with experimental observations that collective dissociation is favored in moderately adhesive environments, where leader cells generate traction while followers maintain partial intercellular bonds [30, 32]. The observed enrichment of leader-dominated, intermediate-sized clusters parallels partial epithelial–mesenchymal transition (pEMT), in which cells retain adhesive contacts while acquiring motility. Notably, the cluster size range observed here (2–50 cells) overlaps reported sizes of CTC clusters with higher metastatic efficiency relative to solitary cells [11, 29]. Together, these findings provide a mechanistic explanation for why intermediate adhesion and strong migration create a regime that fosters cluster detachment, leader-driven persistence, and ultimately, metastatic dissemination.

### Proliferation Plays a Limited Role in Tumor Invasion Dynamics

We next examined the extent to which follower proliferative probability (PP) regulates invasion behavior relative to adhesion and migration. For each simulation (13,310 total), we systematically perturbed a single parameter while fixing the other two, allowing us to quantify marginal effects on invasion metrics and phenotype classification across the full parameter cube.

Across all five invasion metrics, PP showed only minimal influence compared with the migration coefficient and contact energy (Fig 4A–E). Increasing the migration coefficient (λ) consistently elevated the invasive and infiltrative area, promoted the formation of fingers, and sharply increased the number of singles and detached clusters. Contact energy (*J*_*lf*_) exerted a non-monotonic effect, with peaks at intermediate adhesion (*J*_*lf*_ ≈ 0–2) that supported cluster formation and cohesive protrusions. By contrast, proliferative probability produced nearly flat curves, with only minor oscillatory deviations. The most consistent effect of PP was on the overall invasive area, which increased under high proliferation due to expansion of the tumor core, rather than changes in invasive mode. Importantly, shaded bands denote mean *±* SEM across binned parameter values, confirming robustness of these observations.

**Fig 4.**
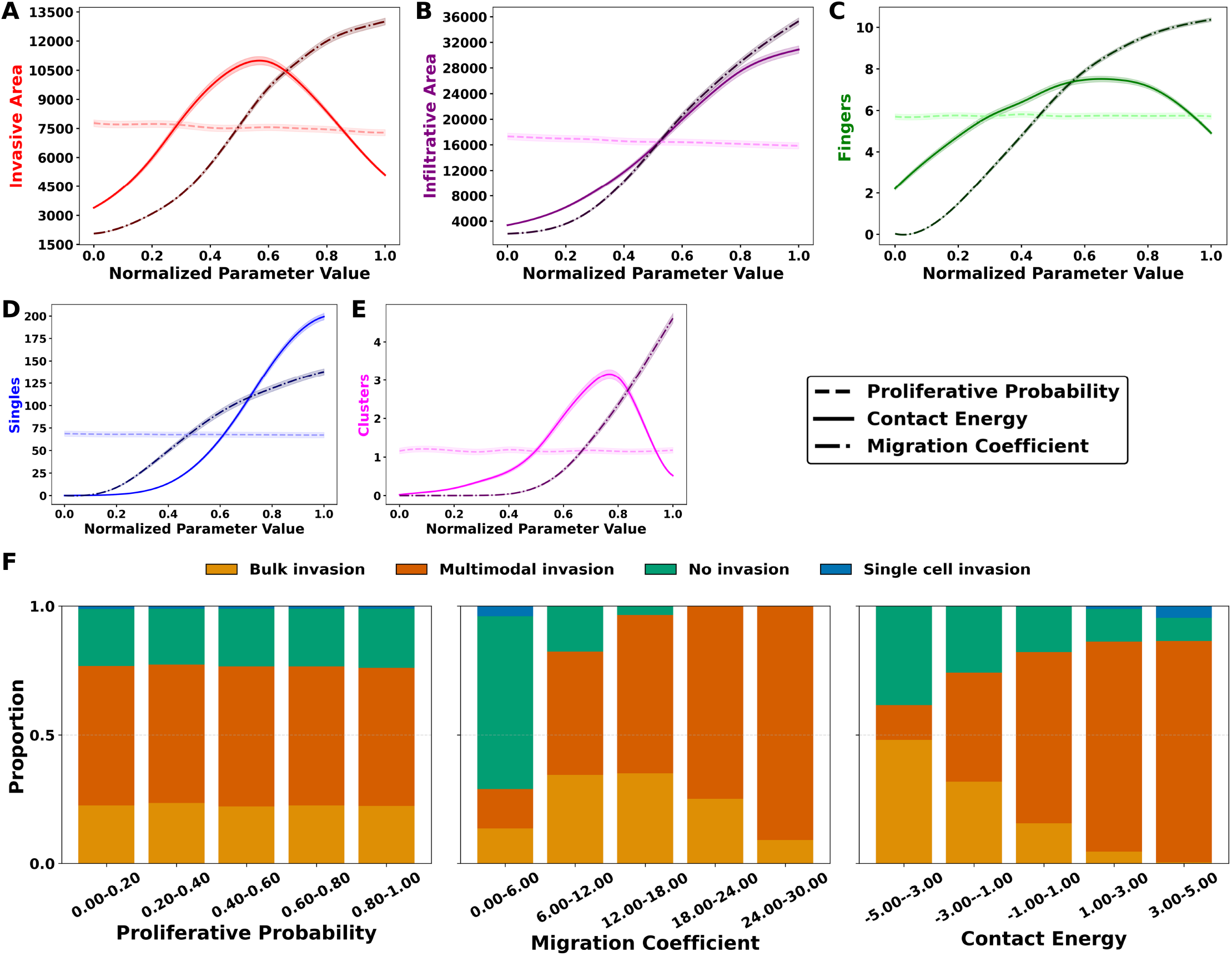
Proliferation minimally affects invasion compared with migration and adhesion. (**A–E**) Metric-parameter relationships for invasive area (A), infiltrative area (B), fingers (C), singles (D), and detached clusters (E). X-axis is a normalized parameter value (scaled to [0, 1]). Original parameter ranges were *λ* ∈ [0, 30] (migration coefficient) and *J*_*lf*_ ∈ [−5, 5] (contact energy, lower values = stronger adhesion). Curves show marginal dependence on proliferative probability (dashed, light shade), contact energy (solid, base shade), and migration coefficient (dash-dot, dark shade). Shaded bands denote mean *±* standard deviation. Migration strongly increased invasive and infiltrative areas, singles, and clusters. Contact energy shows non-monotonic optima at intermediate adhesion, and proliferation induces minor changes. (**F**) Phenotype composition across PP bins remains largely unchanged, whereas migration and adhesion drive substantial phenotype transitions.

Phenotype classification mirrored these trends (Fig 4F). The proportion of tumors across bulk, multimodal, single-cell, and non-invasive classes remained effectively unchanged across proliferative probability (PP) bins. In sharp contrast, both λ and *J*_*lf*_ produced marked phenotype transitions (Fig 4F, inset).

Statistical sensitivity analyses reinforced these results. Distance correlation analysis (S7A Fig) revealed strong dependencies of invasion metrics on migration (up to 0.79 for fingers) and moderate dependencies on adhesion (e.g., 0.68 for singles), while PP correlations were negligible (≤ 0.02). A multi-output random forest model trained on all three parameters (400 trees, mean-decrease-in-impurity criterion) showed a similar pattern (S7B Fig): migration (~ 0.51) and adhesion (~ 0.49) accounted for virtually all explanatory power, while proliferation contributed negligibly (~ 0.01). Pearson and Spearman correlation coefficients similarly showed near-zero correlations between PP and all metrics (|r| < 0.03).

Together, these results establish that proliferation affects the scale of tumor growth, reflected in a higher invasive area at high PP due to an expanded core, but it does not alter the structure or mode of invasion. Migration and adhesion remain the primary regulators of spatial architecture and invasion mode, while proliferation contributes only to tumor mass expansion.

### Discussion

Collective invasion in tumors emerges from a small set of mechanical ingredients. In this work, we focused on three: cell–cell adhesion, directed leader motility, and follower proliferation. Within a leader-follower Cellular Potts framework, these parameters are encoded as leader-follower contact energy *J*_*lf*_ [25, 33], migration coefficient (λ), and a proliferation probability PP for followers.

The leader-follower adhesion is plausibly tuned by cadherin balance: E-cadherin strengthens epithelial junctions, while N-cadherin weakens epithelial cohesion and couples to pro-migratory signaling [34, 35]. The migration coefficient (λ) summarizes the leaders’ responsiveness to external directional cues, regardless of the guidance mechanism. In vivo, invasive fronts likely integrate multiple taxis, including chemotaxis toward soluble gradients, haptotaxis along matrix-bound ligands, and durotaxis up stiffness gradients [36–38]. Therefore, we view λ as an effective directional sensitivity of leader cells. Proliferation represents growth-driven expansion and crowding.

Across 13310 simulations, we consistently observed four qualitatively distinct invasion pehnotypes: No Invasion, Bulk Collective, Single-Cell, and a dominant Multimodal state. This observation parallels the conceptual invasion spectrum proposed by Friedl et al. [5]. Invasion mode is governed primarily by the nonlinear coupling between adhesion and motility, with proliferation acting mainly as a scaling factor for tumor bulk. Multimodal invasion, characterized by coexisting cohesive fingers, solitary cells, and detached clusters, occupies a broad regime of intermediate adhesion and high leader motility. In this region, leader traction is strong enough to deform and partially break cohesion, but adhesion is not so weak that all cells disperse as single defectors. At very weak cohesion, invasion shifts toward predominantly single-cell escape; at strong cohesion and sufficiently high motility, invasion remains collective and compact.

These phase-space patterns connect naturally to experimental observations of leader–follo systems. Leader/follower subpopulations identified in SaGA-profiled NSCLC spheroids often exhibit mixed invasion with coexisting strands, clusters, and solitary cells [12, 17]. Our phase diagram suggests that such behaviors correspond to an intermediate adhesion and high motility regime ((*J*_*lf*_ ~ 1–3) λ ~ 20–30). Interventions that stabilize adherens junctions (e.g., overexpressing α-catenin [39] or strengthening E-cadherin function) would effectively decrease *J*_*lf*_, suppress cluster detachment, and push tumors toward bulk collective, a prediction testable through CTC count in blood. Conversely, tumors exhibiting purely single-cell invasion under very weak adhesion might be steered first into a more cohesive bulk state by pharmacologically reinforcing adhesion, creating an intermediate state where subsequent motility inhibition can arrest invasion.

The multimodal phenotype is consistent with partial epithelial–mesenchymal transition (pEMT) [32, 40–42], where cells co-express epithelial and mesenchymal markers and generate a mix of collective and solitary behaviors. Our model shows that a similar continuum of invasion behaviors arises purely from the mechanical balance of adhesion and motility, without explicitly encoding intermediate transcriptional states. In that sense, pEMT-like behavior in our simulations is an emergent property of effective mechanical parameters. Coupling the CPM mechanics to gene regulatory models of EMT would allow direct comparison between EMT state dynamics and the mechanical phase diagram: for example, testing whether multi-stable EMT circuits sharpen the boundaries between bulk, multimodal, and single-cell regions or simply bias the system toward particular parts of the parameter space.

A consistent outcome of our analysis is that proliferation controls tumor mass but not invasion architecture. This separation between growth control and structural control has practical implications. Anti-proliferative agents are expected to reduce total metastatic burden by decreasing the number of disseminating cells, but the model suggests they are unlikely on their own to convert a multimodal, cluster forming invasion front into a compact, non-invasive growth. In contrast, drugs that target adhesion or motility map onto *J*_*lf*_ and λ and are predicted to reconfigure the invasion architecture itself [43, 44]. Combining these two classes of interventions, one that reshapes invasion patterns and on that scales cell numbers, provides a mechanistic rationale for experimental strategies aimed at reducing metastatic risk.

Our phase diagram can thus be viewed as a control surface for steering tumors between invasion modes. Sensitivity analysis identifies leader motility as the dominant driver of dispersal and intermediate-to-strong adhesion as a consolidating force. For tumors exhibiting multimodal invasion, inhibiting leader motility, for instance, through targeting Rac, Rho/ROCK, or focal adhesion kinase (FAK) pathways with agents such as defactinib or VS-6063 [43, 45], should move the system leftward in the phase space by lowering λ. Agents that modestly increase E-cadherin function or stabilize adherens junctions, such as compounds that enhance α-catenin recruitment [39, 46], would lower *J*_*lf*_, tethering potential defectors and clusters to the primary mass. For tumors invading primarily as single cells under very weak cohesion (*J*_*lf*_ > 3), adhesion reinforcement is the critical first step, even if it transiently promotes bulk collective invasion, because it creates a state where subsequent motility inhibition can arrest invasion.

The parameters *J*_*lf*_, λ, and PP can also be interpreted in terms of measurable biomarkers. Low *J*_*lf*_ corresponds to strong leader–follower adhesion and correlates with high E-cadherin and membrane-localized β-catenin, while high *J*_*lf*_ corresponds to weak cohesion and correlates with cadherin switching, vimentin upregulation, or nuclear β-catenin [35, 47]. The migration coefficient λ reflects the integrated effect of chemokine receptors (e.g., CXCR4), integrin-mediated adhesion to matrix, and intracellular polarity circuits. It could be estimated from organoid invasion assays by measuring leader velocity and directional persistence [48, 49]. Proliferative probability PP corresponds to standard proliferation markers such as Ki67 or cell cycle reporters [50].

Several simplifications in the current model point to natural extensions. First, the 2D slab geometry collapses the full 3D structure of spheroids and tissue. Preliminary 3D simulations (S1 Fig, S3 Video) suggest that qualitative features such as finger formation and cluster detachment are preserved, but the parameter thresholds for intermediate adhesion window may shift, and new escape routes may emerge along the third dimension. Second, the extracellular matrix is static and homogeneous. In reality, cell migration remodels collagen, align fibers, and carve guidance tracks that can feed back on motility and cohesion [51–53]. Including a dynamic fiber network and explicit matrix mechanics whould clarify how ECM remodeling reshapes the boundaries between bulk, multimodal, and single-cell regimes. Third, proliferation is treated as spatially uniform and leader/follower identities are fixed. In vivo, cell-cycle status and EMT state are spatially heterogeneous and plastic on timescales comparable to our simulations. Allowing stochastic switching between leader and follower phenotypes, and imposing nutrient or oxygen gradients that modulate PP could reveal whether plasticity stabilizes multimodal invasion or induces temporal switching between invasion modes.

Finally, this model does not include explicit immune or stromal components. Tumor-associated macrophages, cancer associated fibroblasts, and cytotoxic lymphocytes all reshape the effective values of *J*_*lf*_ and λ by remodeling ECM, secreting soluble factors, and exerting selective pressure on different invasion strategies [10, 54,55]. For example, strong immune predation on solitary cells may indirectly favor cluster-based dissemination, consistent with the high metastatic efficiency of CTC clusters [29, 30]. Embedding our leader–follower mechanics into a richer microenvironment with stromal and immune agents is therefore a natural next step that may shift or fragment the current phase boundaries.

Despite these simplifications, the present work provides a high-resolution, mechanistically interpretable map linking cell-level adhesion and motility to tissue-level invasion strategies. The prevalence of multimodal invasion in our simulations argues against a strictly binary view of collective versus solitary invasion and instead supports a spectrum in which clusters, fingers, and single cells coexist over a wide range of conditions. By making the model and analysis code available and framing invasion control as movement on a phase diagram, we also aim to provide a reusable computational tool for designing experiments that shift tumors away from cluster-forming, metastasis-prone states by jointly modulating cohesion and motility.

### Model and Methods

#### Computational Model of Cancer Invasion

To study the collective invasion of solid tumors, we developed an in silico model using the software CompuCell3D (CompuCell3D.org) version 4.6.0 [56], a modeling environment implementing the lattice-based Cellular Potts Model (CPM) framework. The CPM is a stochastic, energy-minimizing framework for simulating multicellular systems in which cells are represented as extended, deformable domains on a discrete lattice. The mathematical formulation of the CPM energy functional and the Metropolis algorithm governing system dynamics are detailed in S1 Appendix.

The computational tumor represents key features of non-small cell lung cancer (NSCLC) tumor spheroids used in experimental systems such as the Spatiotemporal Genomic and Cellular Analysis (SaGA) platform [12]. While tumor invasion typically begins as 3D spheroids embedded in extracellular matrix (ECM), we simplified this geometry using a quasi-two-dimensional cross-section, discretized on a 500 × 300 lattice with periodic boundary conditions along the horizontal (x) direction, as illustrated in Fig 5. This domain size was chosen to accommodate invasive fronts extending up to 200 pixels from the initial tumor while maintaining computational tractability for large-scale parameter sweeps.

**Fig 5.**
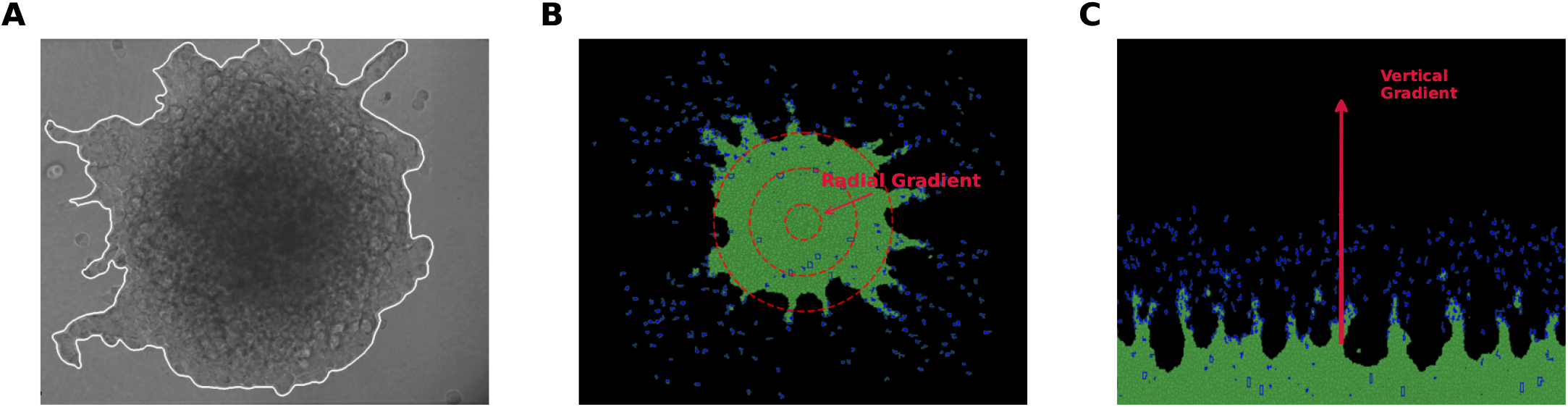
Geometry abstraction from experimental spheroids to in silico domains. **(A)** Experimental tumor spheroid embedded in ECM with leader-driven finger-like protrusions at the invasive front. **(B)** In silico spheroid under a radial migratory field; dashed concentric rings indicate radial distances, and the “Radial Gradient” annotation marks outward bias. **(C)** Rectangular slab abstraction used in this study. Cells are initialized at the base and exposed to a vertical stimulus that increases along the y-axis (arrow). This reduced 2D geometry preserves directional invasion behavior while enabling reproducible, high-throughput parameter sweeps. S3 video illustrates the temporal evolution of the corresponding 3D simulation, confirming that the reduced 2D slab abstraction effectively captures the key structural features of tumor invasion.

In adopting this 2D abstraction, we made three simplifying assumptions, each supported by prior computational work or our own preliminary analyses. First, 2D simulations can qualitatively reproduce key 3D invasion dynamics. Several prior CPM studies [57–59] have demonstrated that 2D cross-sections preserve essential morphological features such as finger formation, growth dynamics, and spatial patterning observed in 3D tumor models. Our preliminary 3D spheroid simulations (S1 Fig, S3 Video) confirmed that invasive area, finger counts, and cluster incidence differed by less than 15% between 2D cross-sections and full 3D simulations under equivalent parameter sets, validating our geometric abstraction for the purposes of phenotype mapping.

Second, ECM remodeling does not qualitatively alter invasion modes within the simulation timeframe. We have previously quantified collagen fiber alignment as a consequence of cancer cell invasion over 24-hour periods [52, 53], finding that significant matrix reorganization occurs on timescales of 24–48 hours. Since our simulations span approximately 36 hours (700 MCS), the static ECM assumption is reasonable for capturing initial invasion dynamics before substantial matrix feedback becomes dominant. Third, the tumor front curvature inherent to spheroid geometry does not fundamentally change the parameter dependence of invasion phenotypes. While absolute metric values may shift between planar and curved fronts, the transitions between no invasion, single-cell, bulk collective, and multimodal invasion are governed primarily by the balance of adhesion and motility, which are preserved in the 2D slab geometry.

We initialized the simulated tumor as a planar slab of follower cells (FCs) occupying the bottom of the domain, spanning the full width (500 pixels) and a height of 21 pixels (y = 0 to 21), corresponding to approximately 52.5 µm in biological units (see Spatiotemporal Calibration). This configuration contained approximately 1167 follower cells, with exact counts varying slightly due to stochastic initialization. A random subset of 25% of these cells was then reassigned as leader cells (LCs), consistent with experimental observations of leader fractions around 30% in NSCLC spheroids [54]. This configuration mimics the basal cross-section of a spheroid tumor embedded in ECM, maintaining invasion polarity and architecture observed in vitro.

A spatially static, non-diffusing chemoattractant gradient field 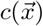 (implemented as a CompuCell3D chemical species named ‘MV’ for migration vector) was imposed along the y-axis to simulate a constant directional migration cue. The gradient was defined as c(x, y) = y/g, where g = 1 sets the gradient strength. This linear gradient represents a fixed directional stimulus consistent with vertical invasion observed in Matrigel-based assays [60], and could correspond biologically to oxygen gradients, growth factor gradients, or other chemotactic cues that guide leader cell migration away from the tumor core. This quasi-2D slab configuration preserves essential directional invasion dynamics while reducing computational complexity, allowing for systematic exploration of key biophysical parameters across more than 13,000 simulations.

### Spatiotemporal Calibration

The correspondence between simulation spatiotemporal scales and in vitro tumor cell behavior was established by calibrating leader cell migration speeds to experimentally measured values from 3D collagen-based invasion assays [12]. Leader cells in these assays exhibited mean velocities of (1.1 ± 0.3) µm/min based on manual tracking of individual cells over 6-hour time windows.

To determine the spatial and temporal conversion factors, we first measured leader cell displacement in our simulations at λ = 20 (intermediate motility) over 100 MCS windows, obtaining an average displacement of 0.4 px/MCS. We set the spatial scale to 1 px ≈ 2.5 µm, guided by typical carcinoma cell diameters of 15 µm to 20 µm. The target cell volume of 10 px^2^ corresponds to a circular cell with diameter 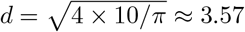 pixels ≈ 8.9 µm, consistent with the cross-sectional area of NSCLC cells in 3D culture. This spatial scale yields a target cell area of 10 px^2^ ≈ 62.5 µm^2^.

With this spatial scale, the observed displacement of 0.4 px/MCS corresponds to 1.0 µm/MCS. Based on calibration against the experimental invasion time course and cell cycle dynamics, we set the temporal scale to 1 MCS ≈ 3.1 min, yielding simulated leader velocities of approximately 1.0 µm/3.1 min = 0.32 µm/min at λ = 20. For higher migration coefficients (λ ≈ 30), simulated velocities approached 0.9 µm/min to 1.2 µm/min, matching the range of experimental measurements.

This spatiotemporal conversion factor (1 px ≈ 2.5 µm, 1 MCS ≈ 3.1 min) was adopted for all analyses. Accordingly, our standard simulation duration of 700 MCS represents approximately 36 hours of biological time, consistent with the 24–48 hour timeframe of SaGA-based invasion imaging [12, 17]

### Simulation Parameters and Parameter Scan Design

The model consisted of two biologically distinct cell types: leader cells (LCs), which exhibit strong directed migration, and follower cells (FCs), which proliferate stochastically but lack intrinsic motility. Leader cells made up a fixed fraction (25%) of the initial tumor and were stochastically reassigned from the follower pool at initialization, as described above. We note that changing the initial leader fraction would likely influence invasion modes and the resulting phase diagram; systematic exploration of this parameter is reserved for future work.

Volume constraints were imposed on all cells, with a target volume V_0_ = 10 lattice units and volume constraint strength λ_V_ = 2.0. The target volume of 10 pixels corresponds to approximately 62.5 µm^2^ cross-sectional area, consistent with a spherical carcinoma cell of 10 µm diameter. The volume constraint strength λ_V_ = 2.0 was chosen to allow realistic cell shape fluctuations while preventing unrealistic cell fragmentation or excessive volume deviation, following previous CPM tumor models [57]. This value balances mechanical stability with biological shape variability observed in time-lapse microscopy of migrating cancer cells.

The main simulation study examined the impact of three biophysical control parameters on tumor invasion dynamics. The first parameter, leader–follower contact energy (*J*_*lf*_), regulated the adhesive interaction strength between LCs and FCs and was varied across a range of −5 to +5 in 11 discrete levels. In the CPM framework, higher contact energy (*J*_*lf*_) corresponds to weaker adhesion between cell types, while lower (more negative) *J*_*lf*_ indicates stronger adhesive interactions. The second parameter, migration coefficient (λ), modulated the responsiveness of LCs to the chemoattractant gradient 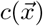 ranging from 0 to 30 in 11 discrete levels. This parameter represents the migratory sensitivity and can be interpreted as the effective force per unit concentration gradient experienced by leader cells. The third parameter, proliferative probability (PP), represented the fraction of follower cells that are proliferation-competent, with values ranging from 0 to 1.0 in 11 discrete levels.

These three parameters were explored over 11 discrete levels each, chosen to balance parameter space resolution with computational cost, resulting in a total of 11^3^ = 1,331 unique parameter combinations. To address the inherent stochasticity of the Cellular Potts Model and ensure statistical robustness, 10 independent replicates were simulated for each parameter set, yielding a total of 13.310 simulations. Each replicate was initialized with a unique random seed (seeds 0–9) to sample different stochastic trajectories.

These parameter ranges were identified through preliminary exploration designed to span the full spectrum of qualitatively distinct invasion phenotypes. Preliminary simulations (n = 50 across broader ranges: *J*_*lf*_ ∈ [− 10, 10], λ ∈ [0, 50], PP ∈ [0, 1]) revealed that parameter values outside the selected ranges produced either uniform behavior or computational artifacts. Specifically, *J*_*lf*_ < −5 resulted in uniformly static, non-invasive tumors due to excessive cell–cell cohesion, while *J*_*lf*_ > 5 caused complete tumor fragmentation into isolated cells. Similarly, λ > 30 induced boundary artifacts due to excessively rapid cell displacement, exceeding the lattice resolution. At the lower end, tumors remained compact with minimal morphological variation. Within the refined ranges (*J*_*lf*_ ∈ [− 5, 5], λ ∈ [0, 30]), we observed transitions among all four invasion phenotypes (no invasion, single-cell, bulk collective, multimodal), confirming that these bounds capture the biophysically relevant parameter space

Follower cell proliferation was implemented using a clock-based system to model asynchronous cell division. At simulation start, a subset of follower cells equal to *PP* × *N*_*FC*_ (where *N*_*FC*_ ≈ 1200 is the initial follower count) was designated as proliferation-competent. Each proliferation-competent cell was assigned an internal mitotic timer, initialized with a random offset uniformly distributed between 0 and 75 MCS to desynchronize division events and avoid artificial synchronization artifacts.

Proliferation-competent follower cells underwent mitosis when two conditions were jointly satisfied; the cell volume exceeded a threshold of 20 px (twice the target volume V_0_, ensuring sufficient mass for viable daughter cells), and when the internal timer surpassed a division threshold drawn from a uniform distribution 𝒰 (25, 125) MCS, centered at 75 MCS. This distribution corresponds to a mean inter division time of 75 MCS ≈ 187.5 min ≈ 3.1 h with a range of approximately 1.0–5.2 hours. This accelerated proliferation rate reflects the rapid growth observed in aggressive cancer cell lines cultured in 3D ECM environments [61], and is consistent with the shortened G1 phase typical of transformed cells. Upon division, the cell was bisected along a randomly oriented axis (CompuCell3D default), and both daughter cells inherited the proliferation-competent state with reset timers drawn independently from 𝒰 (0, 75) MCS, preserving heterogeneity in division timing and spatial organization across the tumor. Cells not initially designated as proliferation-competent remained quiescent throughout the simulation, reflecting the observed heterogeneity in proliferative capacity within tumor populations. Leader cells were not assigned proliferation capacity in the baseline model, consistent with experimental observations that highly motile, invasive leader cells often exhibit reduced proliferation compared to followers [12].

Parameter values, such as proliferation rates, leader cell fractions, and directional motility bias, were motivated by experimental estimates from biological observations on collective cancer invasion [3, 12]. This detailed model setup enabled a robust exploration of how cell-cell adhesion, migration, and proliferation interact to shape the collective invasion behavior of tumors within the CPM framework. A complete summary of all simulation parameters, including dimensions, fixed contact energies, and control parameters, is provided in Table 1.

**Table 1.** Simulation and biophysical parameters used in the leader–follower tumor invasion model. Dimensions are expressed in terms of L (length), T (time), and E (energy). Parameters marked with an asterisk (^∗^) were systematically varied during the parametric scan.

| Parameter | Symbol | Model Value | Dimension |
| --- | --- | --- | --- |
| Boltzmann temperature | $T$ | 10.0 | $E$ |
| Leader cell fraction | $k$ | 25% | – |
| Follower growth rate | $f_{\text{grow}}$ | 0.015 MCS$^{-1}$ | $L^2/T$ |
| Lattice size | – | $500 \times 300$ | $L^2$ |
| Volume constraint strength | $\lambda_V$ | 2.0 | $E/L^2$ |
| Target cell volume | $V_0$ | 10 px | $L^2$ |
| Migration strength* (leaders) | $\lambda$ | 0 – 30 | $E/L$ |
| Proliferative probability* | $PP$ | 0 – 1.0 | – |
| Contact energy*: Leader–Follower | $J_{lf}$ | –5 – +5 | $E$ |
| Contact energy: Medium–Leader | $J_{ml}$ | 2.0 | $E$ |
| Contact energy: Leader–Leader | $J_{ll}$ | 16.0 | $E$ |
| Contact energy: Follower–Follower | $J_{ff}$ | 5.0 | $E$ |
| Contact energy: Medium–Follower | $J_{mf}$ | 10.0 | $E$ |

### Invasion Metrics and Phenotype Classification

For each simulation, spatial and topological features were extracted using a combination of breadth-first search (BFS), neighborhood analysis, morphological data, and cell-type–specific annotation. The invasive area was defined as the cumulative area occupied by the main tumor mass [17]. This was determined by initiating a BFS from the basal layer of tumor cells at the domain floor and identifying all connected cells. This metric reflects collective expansion and requires cells to maintain contact with the core structure. As shown in Fig 6, the invasive front is delineated by the red curve tracing the main tumor boundary. The infiltrative area was computed as the area of the convex hull enclosing all cells and captures the overall spread of the tumor, including disconnected single defectors and clusters [62]. This provides a global measure of spatial dispersion, irrespective of physical connectivity, and is captured in the figure by the orange outermost boundary line.

**Fig 6.**
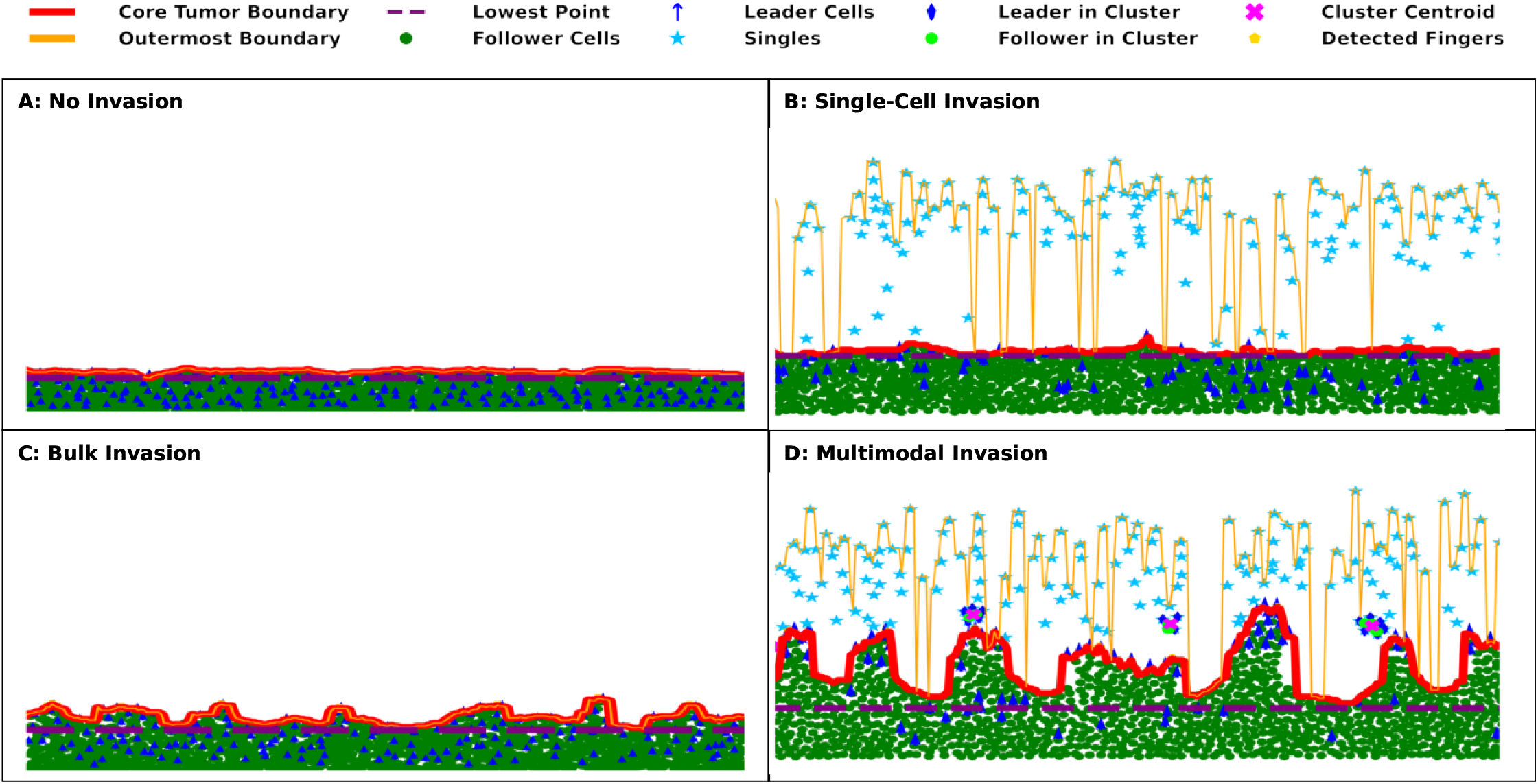
Distinct emergent invasion phenotypes in leader–follower tumor simulations. Representative snapshots of tumor morphologies at endpoint (MCS = 700) for the four canonical invasion phenotypes, each arising from unique combinations of leader motility, follower proliferation, and intercellular adhesion parameters. (**A**) No Invasion: Tumor remains compact with no detachment or dispersal beyond initial bounds. Invasive and infiltrative areas are nearly identical; fingers, defectors, and clusters are absent. (**B**) Single-Cell Invasion: Individual cells (cyan stars) detach from the tumor mass, increasing the infiltrative area without forming collective structures. (**C**) Bulk Invasion: Finger-like protrusions (gold pentagons) emerge, indicative of coordinated collective migration; defectors and clusters are not observed. The invasive area is large and contiguous. (4) Multimodal Invasion: The coexistence of collective fingers, solitary defectors, and detached clusters (magenta Xs) reflects a heterogeneous expansion mode that combines cohesive and dispersed strategies. Standard annotations across panels: core boundary (red), outer boundary (orange), lowest point (purple), follower cells (green circles), leader cells (blue). The legend above the panels illustrates symbol definitions.

For each simulation, spatial and topological features were extracted at the final time point (MCS = 700) using a combination of breadth-first search (BFS), neighborhood analysis, morphological processing, and cell-type–specific annotation. The main tumor mass was identified by constructing a contact graph in which each cell is represented as a node and cell–cell contacts define edges. Starting from all cells at the domain base (y = 1), we performed breadth-first search (BFS) to identify all cells physically connected to the substrate, defining the set of main tumor cells. Cells were considered neighbors if they shared at least one lattice edge (von Neumann neighborhood). This graph-based approach naturally captures the connected component corresponding to the primary tumor while excluding spatially detached clusters and solitary defectors.

The **invasive area** was defined as the cumulative area occupied by the main tumor mass [17], computed by summing the volumes of all cells belonging to the BFS-identified connected component anchored at y = 1. This metric reflects cohesive tumor expansion and requires cells to maintain physical contact with the core structure. The invasive front boundary is illustrated by the red curve in Fig 6. The **infiltrative area** was computed as the area of the convex hull enclosing all tumor cells (both main tumor and detached), capturing the overall spatial footprint of the tumor, including dispersed single cells and clusters [62]. This metric provides a global measure of spatial dispersion, irrespective of physical connectivity, and is represented by the orange outermost boundary in Fig 6.

To characterize morphological complexity, we extracted geometric features from the simulated tumor front. For each x-coordinate, we scanned vertically from the top of the domain (y = 299) downward to identify the highest y-position occupied by a main tumor cell, yielding the tumor boundary profile y_tumor_(x). This one-dimensional boundary was interpolated using cubic smoothing splines (SciPy) to facilitate derivative calculations and peak detection. **Invasive fingers** were detected using a peak-finding algorithm applied to the interpolated boundary profile. Peaks were classified as fingers if they satisfied three criteria; prominence ≥10 pixels in the y-direction (corresponding to ~ 30 µm) protrusion height), separation of at least 20 pixels (~ 60 µm) from neighboring peaks in the x-direction, and width at half-prominence ≥ 5 pixels. These thresholds were chosen to identify biologically meaningful finger-like protrusions while filtering spurious peaks arising from single-cell fluctuations, following established definitions in prior computational studies [26, 27]. To further reduce over-counting of closely spaced peaks, we applied a post-processing merge step: peaks separated by less than 15 pixels were consolidated into a single finger by retaining only the leftmost peak. The resulting count of merged peaks constituted the final finger metric.

**Singles** were defined as individual cells (leader or follower) with no contact neighbors. Specifically, a cell was classified as a single defector if it shared no lattice edges with any other tumor cell (von Neumann neighborhood criterion) and was located above the minimum y-coordinate of the main tumor (to exclude cells at the substrate). This definition captures solitary cells that have fully detached and migrated away from any collective structure. **Detached clusters** were operationally defined as spatially separated groups of two or more tumor cells that maintained internal connectivity but were entirely disconnected from the main tumor component [30]. Clusters were identified using BFS applied to all cells not belonging to the main tumor: for each follower cell outside the main tumor, we initiated a BFS to discover its connected component. If this component contained two or more cells and had no contact with the main tumor, it was classified as a detached cluster. Cluster composition was quantified by counting the number of leaders and followers within each cluster, enabling classification of purely follower-based aggregates, leader-only satellites, and mixed clusters. Cluster centroids were computed as the mean x- and y-coordinates of member cells. Representative examples of annotated features for each invasion phenotype are shown in Fig 6.

Following the computation of invasion metrics, each simulation was assigned a phenotypic label using a rule-based classification framework designed to capture biologically interpretable modes of tumor invasion. This classification scheme utilized a combination of the invasion metrics quantified at the endpoint of each simulation (MCS = 700). The goal was to categorize the emergent invasion behavior into one of four distinct phenotypes: No Invasion, Single-Cell Invasion, Bulk Invasion, and Multimodal Invasion. This classification was designed to reflect experimental categorizations of invasion behavior [5].

Simulations were classified as No Invasion (Fig 6A) if the invasive and infiltrative areas were nearly equal and if there were no fingers, defectors, or detached clusters detected. This indicated that the tumor had remained morphologically compact and did not breach its initial boundaries. The Single-Cell Invasion phenotype (Fig 6B) was assigned to simulations where the infiltrative area exceeded the invasive area, and a non-zero number of solitary defectors were observed, while protrusions and clusters remained absent. This configuration captures the dispersal of individual cells in the absence of coordinated adhesion. Simulations exhibiting a large invasive area accompanied by finger-like protrusions, but no solitary defectors or clusters, were classified as Bulk Invasion (Fig 6C), representing coherent, strand-like collective expansion of the tumor front. Finally, the Multimodal Invasion phenotype (Fig 6D) was assigned when both cohesive structures (e.g., fingers) and dispersed elements (e.g., defectors or clusters) were simultaneously present, and the infiltrative area substantially exceeded the invasive area. This phenotype captures a hybrid mode in which collective and individual invasion coexist within the same spatial domain.

This automated classification pipeline enabled the high-throughput categorization of invasion phenotypes across the whole parameter space of 13,310 simulations. We saved the outputs in CSV format and compiled them into aggregate data tables for phenotype classification. The resulting phenotype labels were subsequently used to construct multidimensional phase diagrams and phenotype frequency distributions, which formed the basis for the phenotypic landscape analysis presented in Fig 2A.

## Data Availability

The computational model is implemented in CompuCell3D, and the source code and simulation data used to produce the simulations and analyses are available in the GitHub repository: https://github.com/Jiang-Lab/Leader_Follower_Invasion_Model.

## Acknowledgments

We thank Somiya Rauf, Sima Moshafi, and Wilbur Hudson for their helpful inputs. Y.J. thanks the Frady Whipple Professorship from Georgia State University.

## Supplementary Information

### S1 Appendix. Cellular Potts Model

All simulations were performed using the open-source simulation environment CompuCell3D [56], based on the Cellular Potts Model (CPM) [24, 25, 63], a lattice-based, energy-minimizing framework for simulating the collective behavior of cells in multi-cellular systems, including cell migration, adhesion, and morphological transitions in tissues. Each cell is represented as a set of connected lattice sites (pixels) sharing a common identifier σ, and interactions between cells evolve by minimizing a global energy functional. The CPM naturally accommodates dynamic changes in shape, adhesion, and movement of cells across space and time.

The total energy of the system is given by:

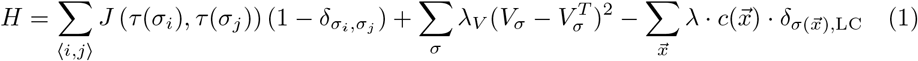

The first term captures cell–cell and cell–medium interactions through a contact energy matrix J(*τ, τ* ^′^), where *τ*(*σ*) denotes the type of the cell occupying lattice site σ. The Kronecker delta function 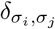 ensures that this energy contribution only arises at interfaces between distinct cells, thus quantifying interfacial tension due to differential adhesion. Lower values of J correspond to stronger adhesion, while higher values indicate weaker adhesive interactions. The second term imposes a volume constraint on each cell. V_σ_ is the current volume of cell σ, and 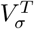 is its prescribed target volume. Deviations from the target are penalized with strength λ_V_, which enforces volume conservation and prevents biologically unrealistic expansion or shrinkage of cells. This constraint is crucial in capturing the physical limitation of space and mechanical pressure within dense tumor tissues. The third term encodes the migratory responsiveness of leader cells. 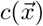 is the local chemoattractant concentration at position 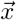 and λ is the migratory sensitivity coefficient that quantifies how strongly leader cells respond to gradients in the chemical field. The Dirac delta function 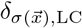 restricts this term to only those lattice sites occupied by leader cells, enabling directed migration along chemoattractant gradients. In this model, follower cells are non-migratory and experience only passive motion through adhesion and growth [12].

Cell configurations evolve stochastically through attempted lattice updates using the Metropolis algorithm, where each attempted cell ID copy from site 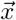 to a neighboring site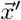 is accepted with a probability governed by the Boltzmann distribution:

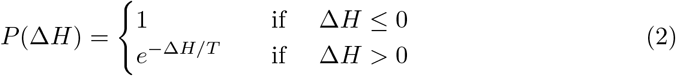

where ∆H is the change in energy associated with a proposed lattice update, and T is the effective simulation temperature controlling membrane fluctuations or biological noise. This framework was implemented using CompuCell3D (version 4.6.0), a modular simulation environment for CPM that incorporates PDE solvers, cell tracking modules, and customizable cell behaviors, such as chemotaxis and volume regulation [56].

### S2 Appendix. Computational Implementation

We use CompuCell3D version 4.6.0 running on Python 3.9. Post-processing used NumPy 1.21, SciPy 1.7, NetworkX 2.6, and Matplotlib 3.4. Each simulation ran for 700 MCS with periodic boundary conditions along the x-axis, requiring approximately 12–18 minutes per replicate on an Intel Xeon E5-2680 processor (2.4 GHz, 8 cores). The complete parameter sweep required approximately 2,800 CPU-hours, distributed across a 48-core computing cluster. Simulations were executed in batches using environment variables to specify parameter combinations (J LF, MU, PP) and replicate indices (REP), enabling efficient parallelization. All simulation code, post-processing scripts, and analysis notebooks are available in the GitHub repository: https://github.com/Jiang-Lab/Leader_Follower_Invasion_Model.

### S3 Appendix. Statistical Analysis

**S1 Fig.**
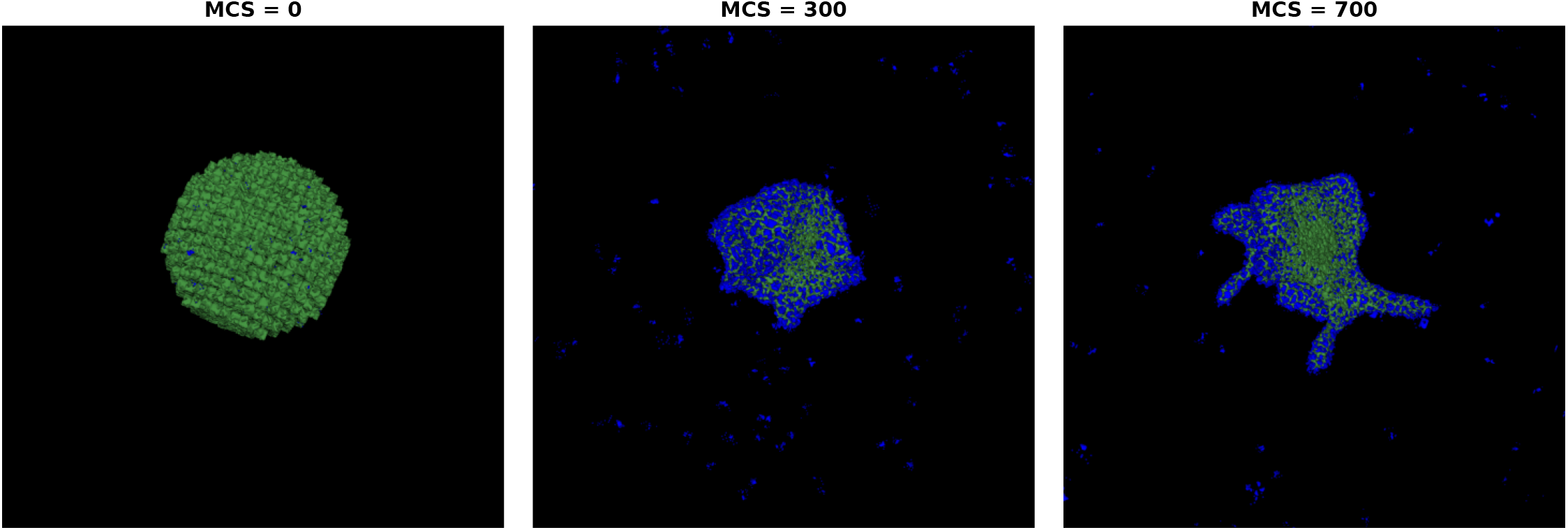
Preliminary simulations in 3D shows 2D domain captures key structural features of tumor invasion. Tumor invasion is visualized at MCS = 0, 300, and 700 using a 3D Cellular Potts Model. The simulation was run with a contact energy between leader and follower cells of *J*_*lf*_ = 2, a motility coefficient of *λ* = 24, and a proliferative probability of *PP* = 0.5. Blue cells represent leader cells, while green cells denote follower cells. These snapshots illustrate the emergence and expansion of invasive structures over time.

**S2 Fig.**
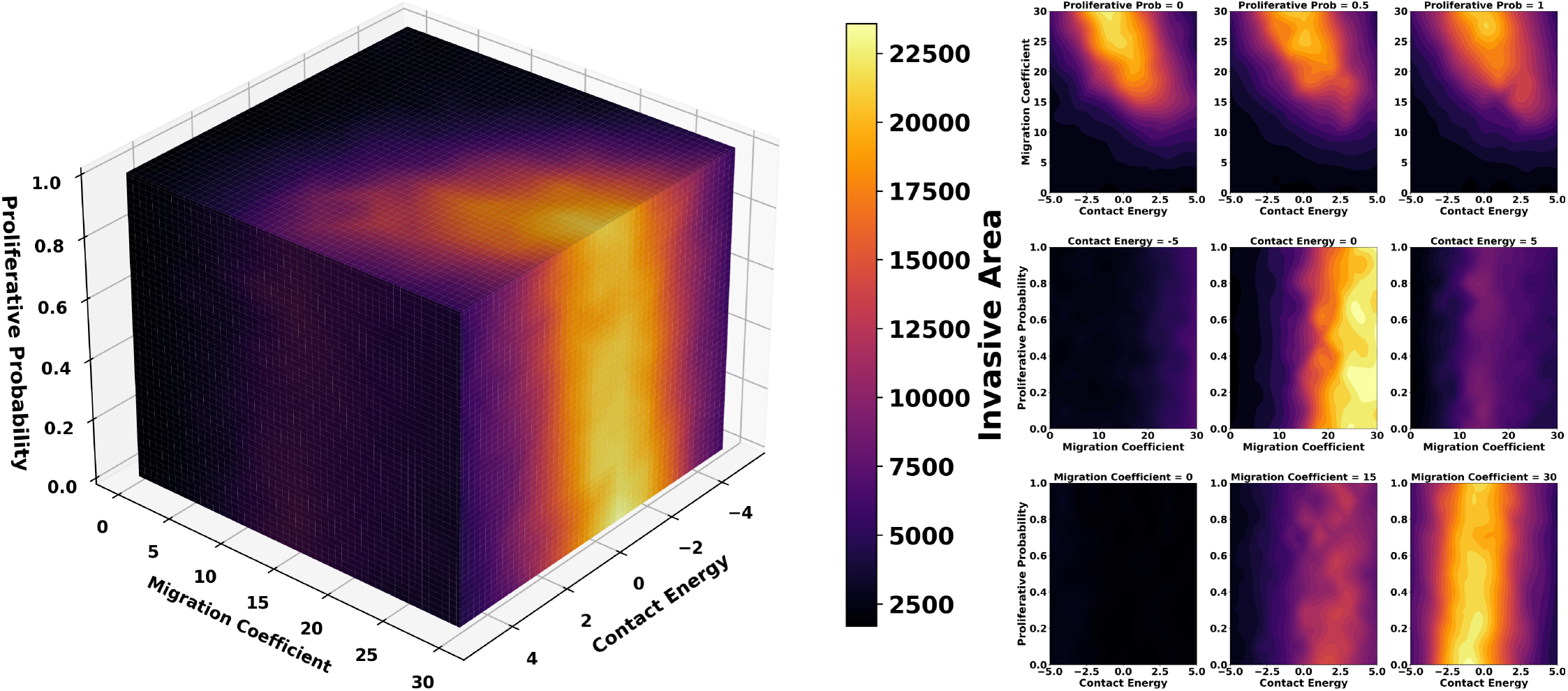
3D volumetric heatmap and cross-sectional views across pairwise parameter slices for the invasive area. Invasive area (tumor spread connected to the primary mass) is shown as a function of leader–follower contact energy (*J*_*lf*_), migration coefficient (*λ*), and proliferative probability (PP). The 3D volume rendering (left) depicts the full parameter space, while 2D slices (right) illustrate representative cross-sections at fixed PP, *J*_*lf*_, or *λ*. Values represent the mean across *n* = 10 replicate simulations for each parameter combination. Shaded gradients denote average tumor size after 700 MCS. Invasive area is maximized under moderate adhesion (*J*_*lf*_ ≈ 0–2) and high migration (*λ* ≥ 15), particularly at elevated proliferation (*PP* ≥ 0.5). Strong adhesion (*J*_*lf*_ *<* − 2) suppresses invasion, while very weak adhesion (*J*_*lf*_ *>* 3) disrupts cohesion, limiting expansion. These results highlight that invasion requires a balance of adhesion and motility, with proliferation modulating overall tumor bulk but not invasion mode.

**S3 Fig.**
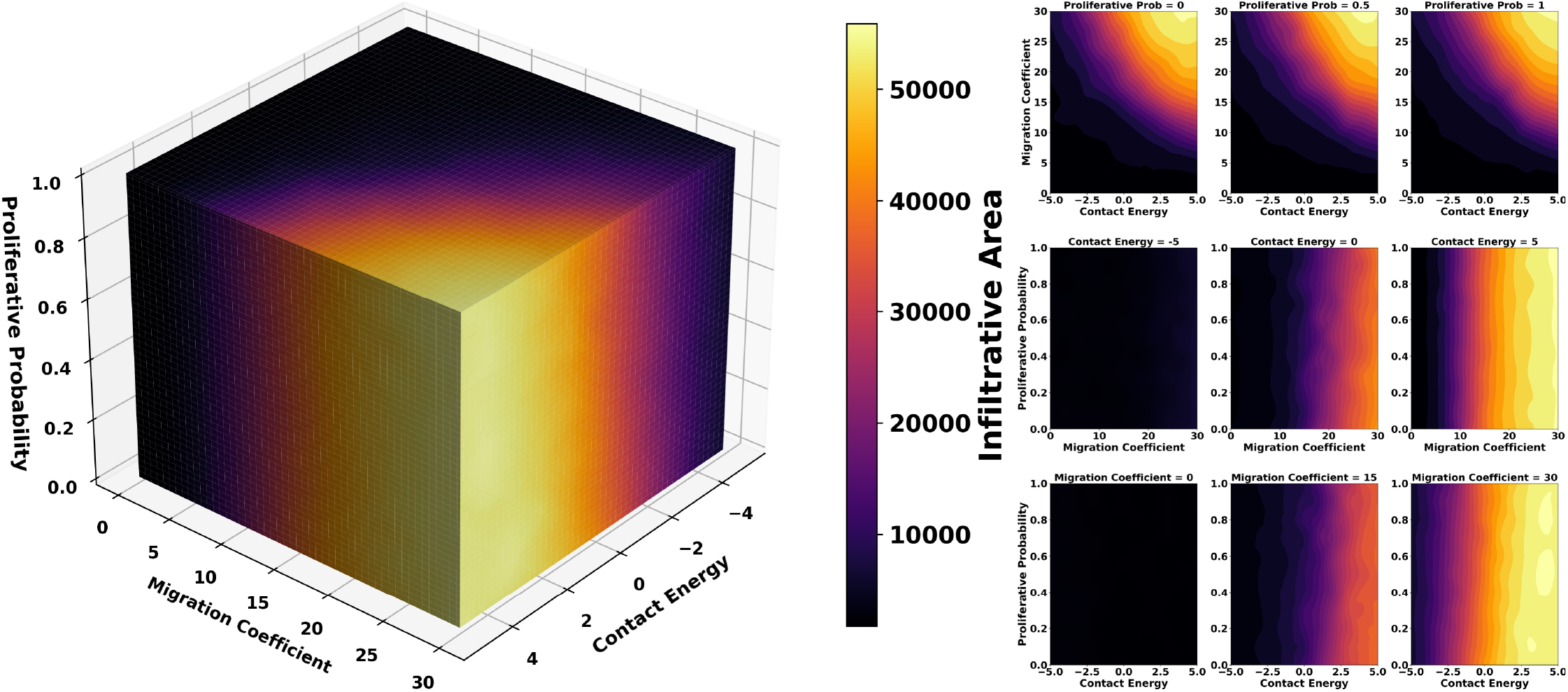
Combined 3D and 2D visualization of the infiltrative area formed across the parameter space. The infiltrative area (global tumor footprint, including dispersed cells and clusters) is mapped across the parameter space. Results are averaged across *n* = 10 replicates per parameter combination. The metric increases monotonically with migration coefficient *λ*, independent of proliferative probability, and peaks under weak adhesion (*J*_*lf*_ *>* 3). Proliferation had minimal impact, as infiltration was driven primarily by motility. Cross-sectional slices show that the infiltrative area expands sharply with *λ* even when PP=0, indicating that leader motility alone is sufficient for extensive spatial spread.

**S4 Fig.**
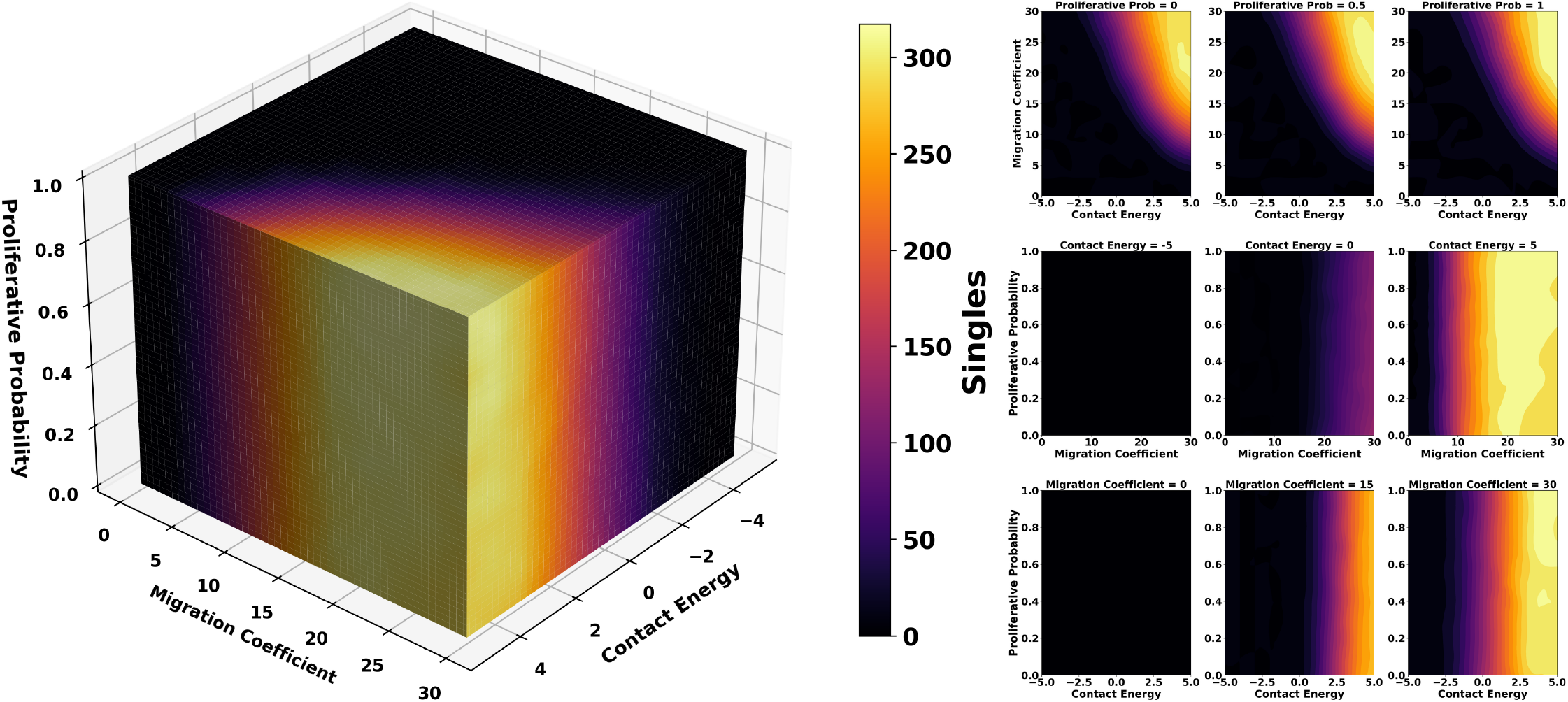
Visualization of isolated cell escape events quantified across parameter slices and in 3D. The number of single-cell defectors (solitary cells that leave the tumor mass) is plotted across the parameter space. Values are means of *n* = 10 replicates. Defectors rise steeply with *λ* under weak adhesion (*J*_*lf*_ *>* 2), reflecting a shift toward mesenchymal-like escape. Proliferative probability (PP) does not significantly affect single defectors, consistent with statistical tests (Pearson | *r* | *<* 0.03). Strong adhesion suppresses solitary escape regardless of motility. These trends indicate that singles arise under regimes of high motility and weak cohesion, independent of proliferation.

**S5 Fig.**
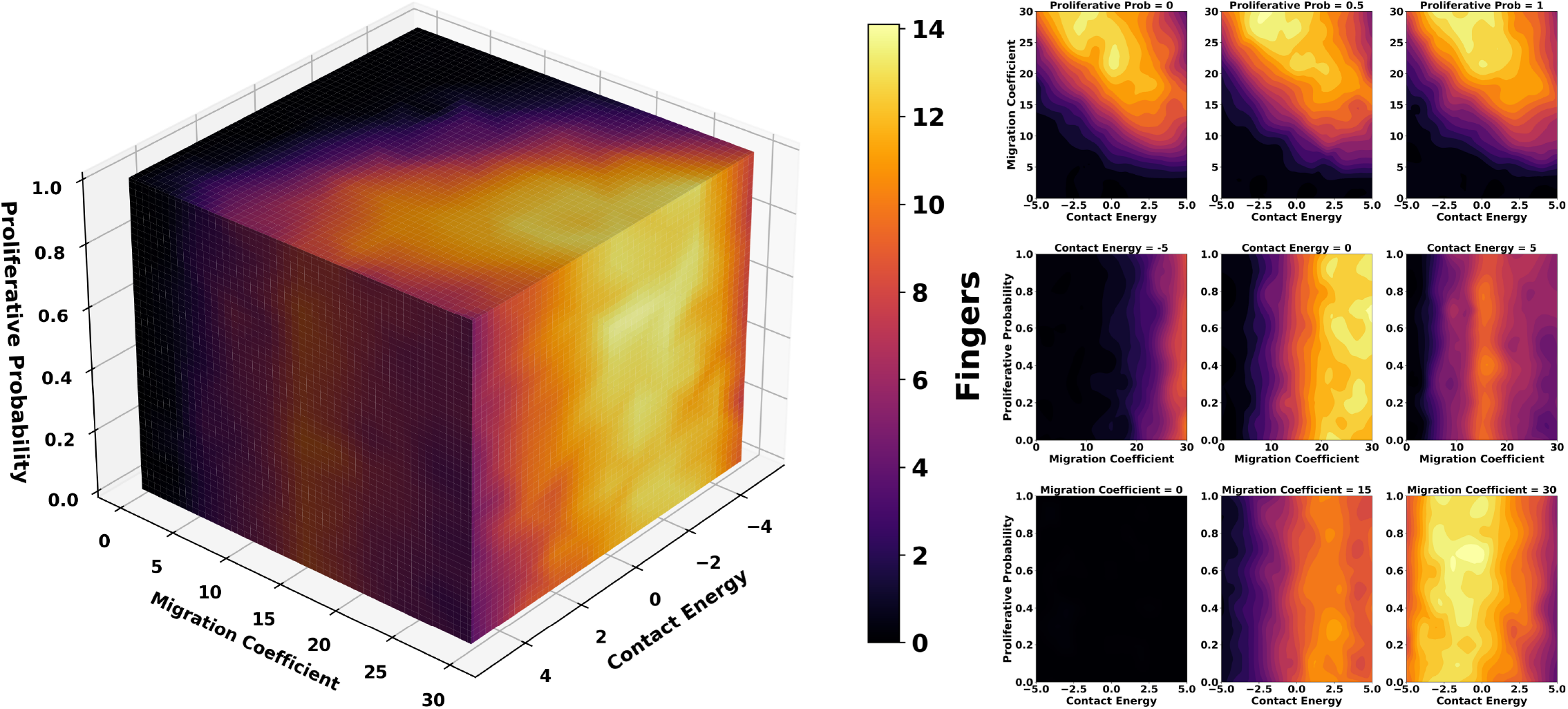
3D Contour plots and cross-sectional views across pairwise parameter slices for the number of protruding fingers. Finger count (cohesive strand-like protrusions) is displayed across migration coefficient, contact energy, and proliferative probability, averaged over *n* = 10 replicates. Finger formation is maximized under intermediate adhesion (*J*_*lf*_ ≈ 0–2) and strong motility (*λ* ≥ 20). Too much adhesion prevents branching, while too little breaks cohesion, leading instead to solitary cells or clusters. PP again had minimal influence on finger incidence. These findings support the existence of an optimal adhesion–motility regime for collective strand-like invasion.

**S6 Fig.**
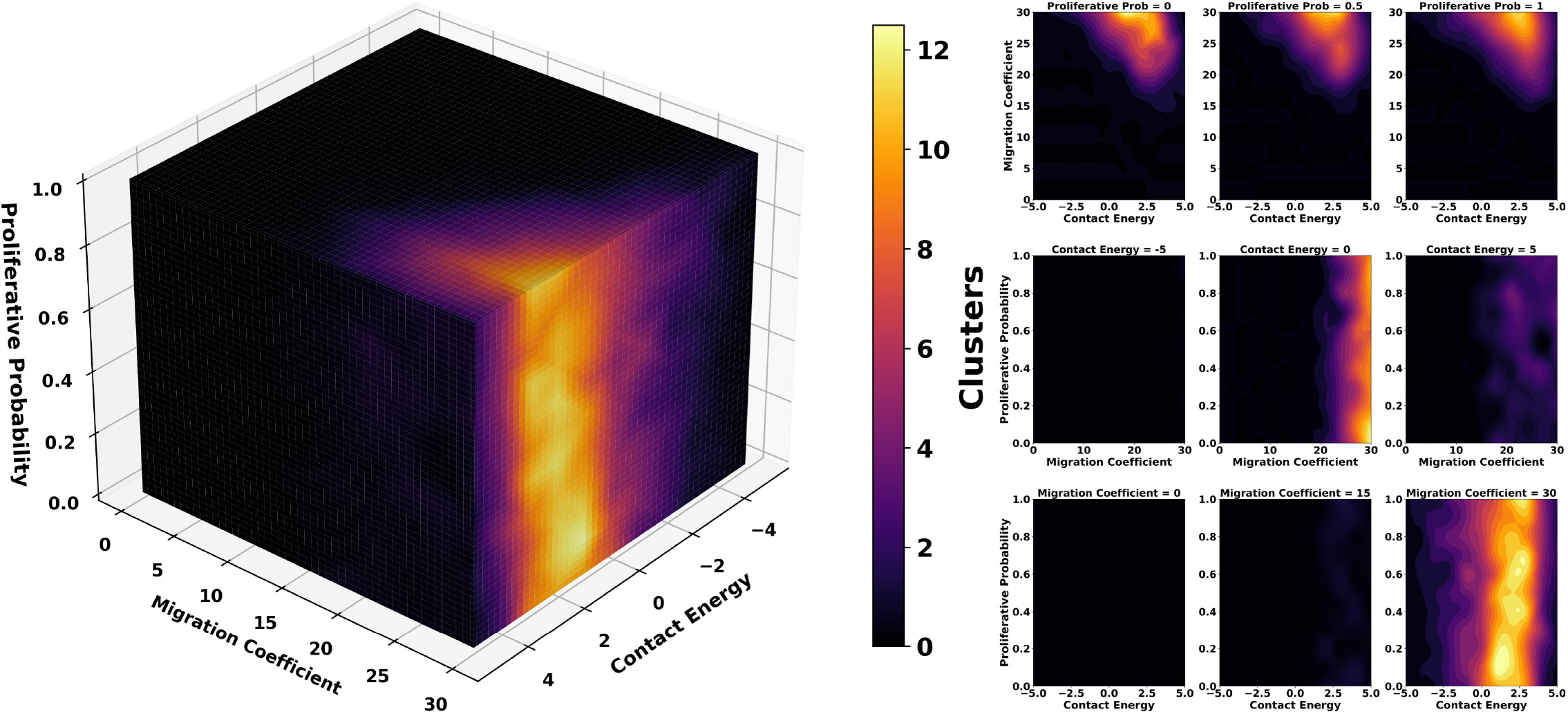
Combined 3D and 2D visualization of the number of clusters formed across the parameter space. Cluster number (detached multicellular aggregates) is quantified across the parameter space. Each value reflects the mean of *n* = 10 replicates. Clusters form predominantly under intermediate adhesion (*J*_*lf*_ ≈ 0–2) and high motility (*λ* ≥ 24), where cohesion is sufficient for cluster integrity but weak enough to allow detachment from the tumor. Proliferation (PP) had a negligible effect on cluster frequency or size. Persistence analysis (not shown) revealed that larger clusters in this regime survived until the end of simulations, whereas smaller aggregates dissolved earlier. These results underscore that cluster-mediated dissemination arises from a narrow adhesion–motility balance rather than proliferative growth.

Parameter sensitivity was quantified using multiple complementary approaches. Pearson and Spearman rank correlations were computed between each parameter (*J*_*lf*_, λ, PP) and each invasion metric (invasive area, infiltrative area, fingers, singles, clusters) to assess linear and monotonic relationships, respectively. Distance correlation was additionally computed to capture nonlinear dependencies not detected by Pearson or Spearman methods.

Feature importance was assessed using a random forest regression model (RandomForestRegressor, scikit-learn 0.24) trained to predict all five invasion metrics simultaneously from the three input parameters. The model consisted of 400 decision trees, and feature importance was quantified as the mean decrease in impurity (Gini importance) averaged across all trees and normalized to sum to 1. This approach provides a model-agnostic measure of each parameter’s contribution to explaining variance in invasion outcomes.

Phase diagrams display the most frequent phenotype across n = 10 replicates per parameter combination. For marginal analyses (Fig 4), parameters were binned into 11 levels corresponding to their discrete scan values, and invasion metrics were averaged within each bin. Error bars and shaded regions represent standard error of the mean (SEM) unless otherwise noted. All statistical analyses were performed in Python 3.9 using NumPy, SciPy, and scikit-learn libraries.

**S7 Fig.**
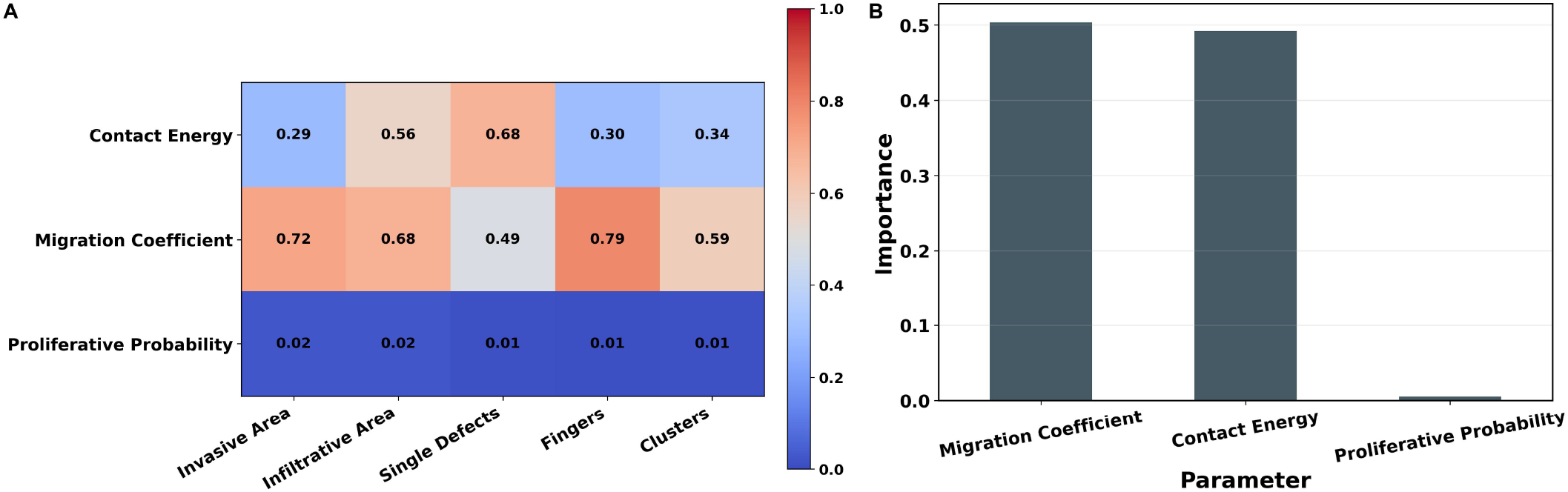
Sensitivity analysis of biophysical parameters with respect to invasion metrics. (**A**) Distance-correlation analysis shows strong dependencies of invasion metrics on migration coefficient and moderate effects of contact energy, whereas proliferative probability remains near zero across all metrics. (**B**) Random-forest feature importance confirms this trend: migration (~ 0.51) and contact energy (~ 0.49) account for nearly all explanatory power, while proliferation (~ 0.01) is negligible. Together, these results demonstrate that invasion phenotypes are governed primarily by motility and adhesion, with proliferation exerting little influence.

**S8 Fig.**
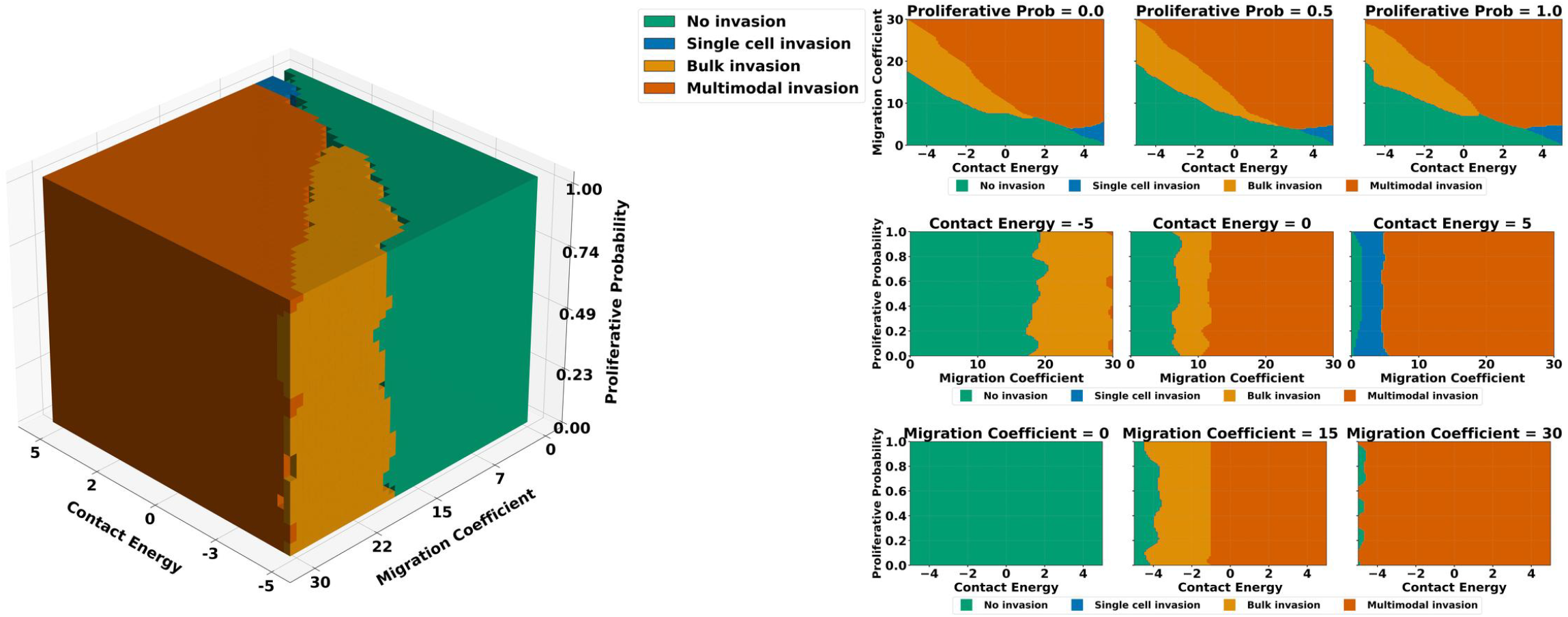
2D cross-sections and orthogonal views of the 3D phenotype phase space. (**A**) The primary 3D phenotype map (reproduced from Fig 2A) visualizes the dominant tumor invasion phenotype at each point in the parameter space defined by contact energy (*J*_*lf*_), migration coefficient (*λ*), and proliferative probability (PP). Phenotypes are color-coded as: green (No Invasion), blue (Single-Cell Invasion), orange (Bulk Invasion), and gold (Multimodal Invasion). (**B**) 2D cross-sections through the 3D phase space clarify the influence of each parameter on invasion outcomes. Horizontal slices (top row) show phenotype distributions across *J*_*lf*_ and *λ* for fixed PP values (0.0, 0.5, 1.0), while vertical slices (middle and bottom rows) show distributions over (*λ*, PP) and (*J*_*lf*_, PP) for selected fixed values of *J*_*lf*_ and *λ*, respectively. These slices reveal sharp phenotype transitions and nonlinear interactions between parameters. The prevalence of multimodal invasion is consistent across a broad range of biophysical conditions.

**S9 Fig.**
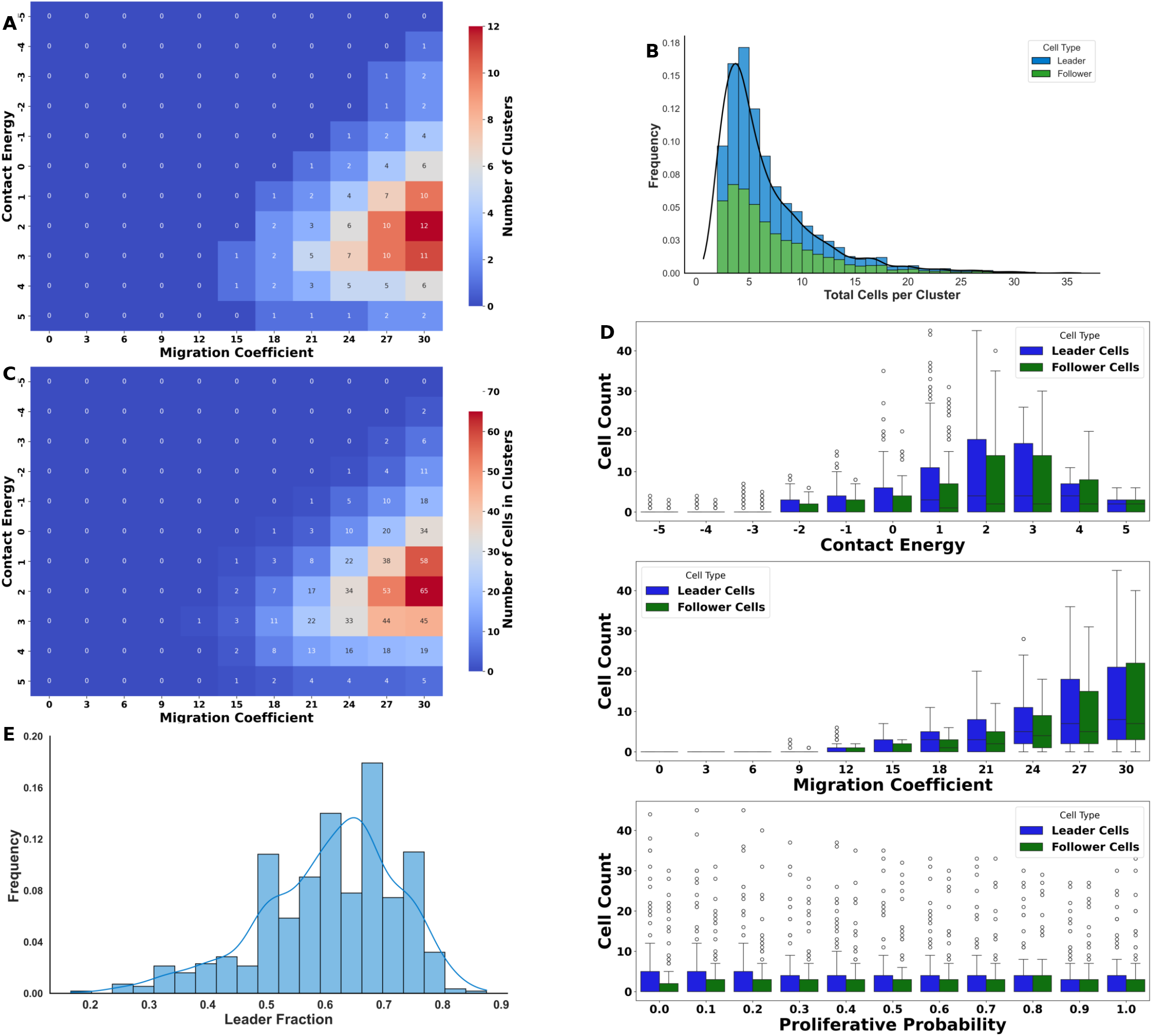
Cluster formation dynamics across adhesion, motility, and proliferation. Summary of cluster-related metrics from 13 310 simulations evaluating the influence of leader–follower contact energy (*J*_*lf*_), leader migration coefficient (*λ*), and follower proliferative probability (PP). (**A**) Heatmap of the average number of clusters across *J*_*lf*_ –*λ* space at fixed proliferation (*PP* = 0.9). Cluster formation is absent in strong-adhesion (*J*_*lf*_ *<* −2) or low-motility (*λ <* 15) regimes, and maximized under intermediate adhesion (*J*_*lf*_ ≈ 0–3) with high motility (*λ* ≈ 24–30). (**B**) Histogram of total cells per cluster, partitioned by leader (blue) and follower (green) identity, reveals a right-skewed distribution: most clusters contain 4–8 cells, while rare aggregates exceed 30 cells. The fitted density curve emphasizes the long-tailed distribution. (**C**) Heatmap of the average number of cells residing in clusters across *J*_*lf*_ –*λ* space, highlighting conditions that yield larger aggregate detachments. (**D**) Box plots of leader (blue) and follower (green) cell counts per cluster under parameter variations. Top: contact energy; middle: migration coefficient; bottom: proliferative probability. Leader cells consistently outnumber followers (~ 4 vs. ~ 3 per cluster on average). Intermediate adhesion and high motility enrich for mixed clusters, whereas increasing proliferation increases follower representation without eliminating leader dominance. (**E**) Distribution of leader fraction (leaders/total cells per cluster), showing a unimodal peak around 0.6. This enrichment indicates that clusters are not random aggregates but leader-dominated groups with robust composition.

### S4 Appendix. Cluster Tracking and Event Detection

Clusters were tracked across timepoints using a centroid-based nearest-neighbor matching algorithm to assign stable identities over the simulation trajectory. At each analysis timepoint (t, sampled every 10 MCS), cluster centroids were computed as the mean (x, y) coordinates of member cells. Clusters at timepoint t + 1 were matched to clusters at t by minimizing centroid displacement, subject to a maximum displacement threshold of 12 px (~ 30 30 µm, corresponding to approximately three cell diameters). Clusters exceeding this displacement threshold were assigned new identities, under the assumption that they represent newly formed aggregates rather than continuations of existing clusters. Splitting events were identified when one cluster at time t matched multiple clusters at time t + 1, indicating fragmentation. Merging events were identified when multiple clusters at t matched a single cluster at t + 1, indicating coalescence. Cluster dissolution was identified when a cluster at time t had no matched descendants at t + 1 and its member cells were either absorbed into the main tumor or dispersed as singles. Overlap fractions for merges and splits were quantified using Jaccard similarity coefficients between member cell sets, thresholded at J ≥ 0.15 to exclude spurious matches. Cluster persistence was defined as the fraction of simulation time (700 MCS) during which a cluster maintained continuous identity through the tracking algorithm.

**S10 Fig.**
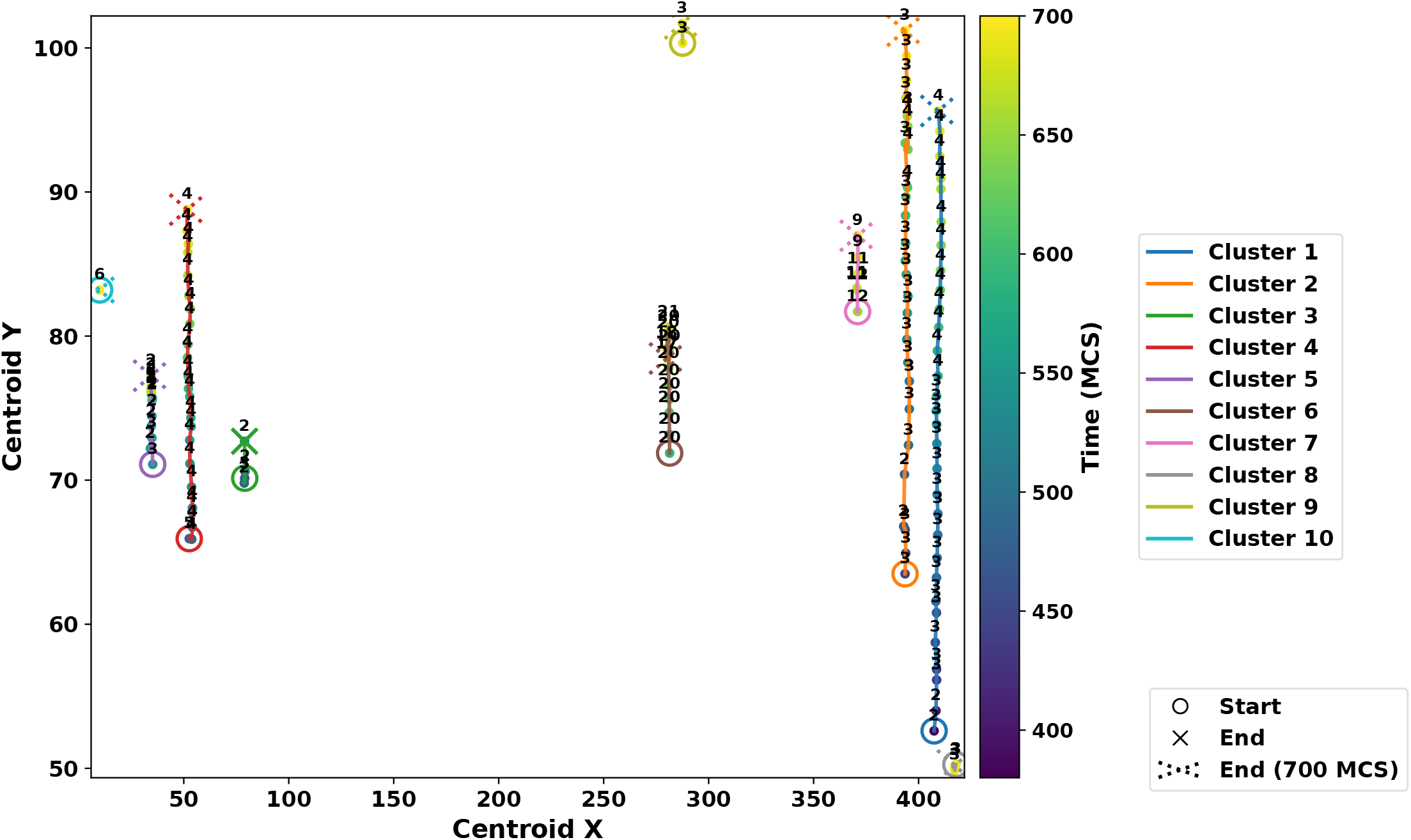
Cluster trajectories with TrackID labeling. Example of cluster centroid trajectories at (*J*_*lf*_ = 1, *µ* = 30, *PP* = 0.9). Each trajectory is colored by cluster identity (TrackID) and annotated with the temporal sequence. Circles denote initial cluster positions. A solid cross marks clusters that terminate before the end of the simulation (disappearance), while a dotted cross marks clusters that persist through the final step at 700 MCS. The color bar encodes simulation time, highlighting splitting, merging, and sustained displacement of leader-enriched clusters. See S2 Video.

**S11 Fig.**
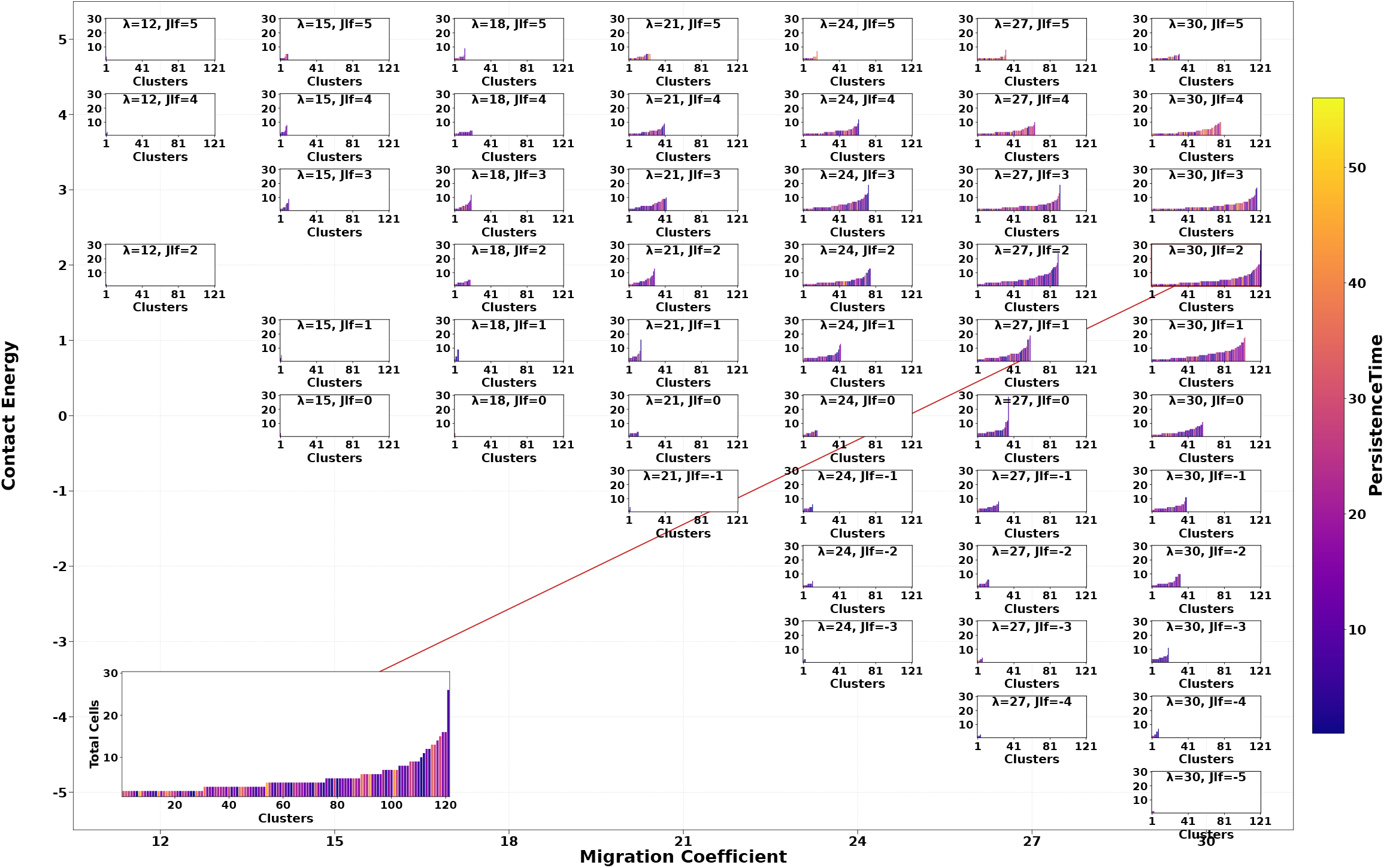
Cluster size distributions across migration and adhesion at fixed proliferation. Shown for *PP* = 0.9 over the (*λ, J*_*lf*_) plane. Each small panel corresponds to one parameter pair and shows the distribution of cluster sizes (y-axis) versus the number of clusters observed (x-axis), pooled across *n* = 10 replicates and sampled every 10 MCS. Bars are colored by persistence time, lighter = longer lived. Panels labeled “No Cluster” indicate no clusters formed under that condition. Cluster formation is absent in strong-adhesion or low-motility regimes, and becomes prominent under intermediate adhesion (*J*_*lf*_ ≈ 0–2) combined with high motility (*λ* ≥ 24). In this window, numerous clusters emerge, typically small to intermediate in size, with occasional large aggregates that exhibit extended persistence. The inset zoom (*λ* = 30, *J*_*lf*_ = 2), displaying the full spectrum of sizes: a predominance of small clusters alongside rare, long-lived large aggregates, illustrating the heterogeneous outcomes characteristic of this parameter regime.

**S1 Table.** Parameter regions associated with emergent invasion phenotypes based on the 3D phenotype map.

| Phenotype | Contact Energy ($J_{lf}$) | Migration Coefficient ($\lambda$) | Proliferative Probability (PP) | Description |
| --- | --- | --- | --- | --- |
| No Invasion | -5 - 2 | 0 - 7 | 0 - 1 | Immotile leaders or excessive cohesion prevent outward expansion. |
| Single-Cell Invasion | 3 - 5 | 4 - 12 | 0 - 0.5 | Weak adhesion and low motility allow leaders to detach and migrate individually. |
| Bulk Invasion | -5 - 0 | 10 - 27 | 0.1 - 0.7 | Strong adhesion and moderate motility facilitate collective strand-like migration. |
| Multimodal Invasion | -1 - 5 | 15 - 30 | 0.3 - 1.0 | Intermediate adhesion with elevated motility and proliferation promotes mixed invasion behaviors. |

**S1 Video. Temporal evolution of Invasion Phenotype Videos** Each panel shows the temporal progression of a distinct invasion mode captured over the course of the simulation. **(A)** Non-invasive phenotype: The tumor remains compact, with no outward expansion, due to strong adhesion, lack of motility, and low proliferation. **(B)** Single-cell invasion: leader cells detach and migrate individually under weak adhesion and high motility. **(C)** Bulk collective invasion: cohesive finger-like projections emerge as a result of strong adhesion, high motility, and active proliferation. **(D)** Multimodal invasion: a heterogeneous mix of strands, solitary cells, and detached clusters forms under moderate adhesion, strong leader motility, and moderate proliferation. Leader and follower cells are shown in blue and green, respectively.

**S2 Video. Dynamic Tracking of Cluster Evolution**. An animation of a representative cluster set is shown over the course of the simulation. The movie highlights key dynamical events, including the splitting of clusters into smaller groups, the merging of detached clusters, and occasional dissolution. These dynamics confirm and complement the static trajectories shown in S10 Fig., particularly illustrating the persistent displacement and interactions of the rightmost clusters.

**S3 Video. Temporal Evolution of Tumor Invasion in 3D Simulation**. Representative movie from the exact simulation shown in S1 Fig. The video spans 0 MCS to 700 MCS, visualizing the spatiotemporal progression of leader (blue) and follower (green) cells. Invasive structures gradually emerge from the spheroid core, with leaders extending protrusions and followers maintaining cohesion. This dynamic view confirms that the 2D projection captures the key structural features observed in the full 3D simulation.

**S4 Video. Temporal evolution of the 3D phenotypic phase space**. Dynamic visualization of invasion phenotype distributions across the parameter space defined by contact energy (*J*_*lf*_), migration coefficient (λ), and proliferative probability (PP), shown from MCS = 0 to 700. Each voxel is colored by the dominant invasion phenotype: green (No Invasion), blue (Single-Cell Invasion), orange (Bulk Invasion), and gold (Multimodal Invasion). Early in the simulations (MCS 0–200), the phase space is dominated mainly by compact, non-invasive tumors. By MCS 300–500, bulk and multimodal invasion emerge under intermediate adhesion and high migration, with isolated regions of single-cell invasion appearing at weak adhesion and high motility. At later stages (MCS 600–700), multimodal invasion becomes the prevailing phenotype across vast regions of the parameter space, confirming it as the dominant strategy. This temporal progression highlights how invasion strategies stabilize over time and demonstrates that multimodal invasion persists as a robust outcome across diverse biophysical regimes.

